# SambaR: an R package for fast, easy and reproducible population-genetic analyses of biallelic SNP datasets

**DOI:** 10.1101/2020.07.23.213793

**Authors:** Menno J. de Jong, Joost F. de Jong, A. Rus Hoelzel, Axel Janke

## Abstract

**Background:** SNP datasets can be used to infer a wealth of information about natural populations, including information about their structure, genetic diversity, and the presence of loci under selection. However, SNP data analysis can be a time-consuming and challenging process, not in the least because at present many different software packages are needed to execute and depict the wide variety of mainstream population-genetic analyses. Here we present SambaR, an integrative and user-friendly R package which automates and simplifies quality control and population-genetic analyses of biallelic SNP datasets. SambaR allows users to perform mainstream population-genetic analyses and to generate a wide variety of ready to publish graphs with a minimum number of commands (less than ten). These wrapper commands call functions of existing packages (including adegenet, ape, LEA, poppr, pcadapt and StAMPP) as well as new tools uniquely implemented in SambaR.

**Results:** We tested SambaR on online available SNP datasets and found that SambaR can process datasets of millions of SNPs and hundreds of individuals within hours, given sufficient computing power. Newly developed tools implemented in SambaR facilitate optimization of filter settings, objective interpretation of ordination analyses, enhance comparability of diversity estimates from reduced representation library SNP datasets, and generate reduced SNP panels and structure-like plots with Bayesian population assignment probabilities.

**Conclusion:** SambaR facilitates rapid population genetic analyses on biallelic SNP datasets by removing three major time sinks: file handling, software learning, and data plotting. In addition, SambaR provides a convenient platform for SNP data storage and management, as well as several new utilities, including guidance in setting appropriate data filters.

**Availability and implementation:** The SambaR source script, manual and example datasets are distributed through GitHub: https://github.com/mennodejong1986/SambaR

## INTRODUCTION

Modern-day population geneticists risk spending as much time studying computer software as studying their actual scientific questions. They also risk spending as much time generating plots as generating new data. These time sinks can negatively affect the quality of research outcomes, as they eat away time needed for a) understanding the theoretical underpinnings of analysis methods and b) interpretation of analysis outcomes.

Integration of computers programs into one single software pipeline removes the necessity of getting acquainted with the technicalities of each program and therefore promotes increased efficiency and, by avoiding incorrect usage, increased accuracy. Efficiency will be improved further if an integrative software package automatically translates the results into ready-to-publish graphs. For two reasons a good candidate for such a wrapper and plotting software is an R package: many tools for population-genetic analyses are written in R (R Core Team, 2019), and R contains powerful graphing tools.

Here we introduce the R package SambaR, which stands for: ‘Snp datA Management and Basic Analyses in R’. SambaR aims to remove the time sinks of a.) managing input and output files, b.) learning computer software and c.) generating and polishing plots. SambaR automates the integrated usage of proven and widely used R packages for population genetic analyses and generates over 100 ready-to-publish graphs to depict data quality control and analyses outcomes.

SambaR is designed to enable population-geneticists to perform a wide variety of population genetic analyses on SNP datasets – quality control, population structure analyses, population differentiation analyses, genetic diversity analyses, and selection analyses – with less than ten commands required. Users are guided through the workflow by an accompanying manual, as well by built-in explicit error messages. By default, SambaR runs most analyses using different methods and/or varying filter and parameter settings, allowing users to explore the data and parameter space and to choose appropriate filter settings.

Apart from streamlining population-genetic analyses, SambaR is also meant to provide a convenient and user-friendly platform for SNP data management. SambaR stores the input data in three data objects. Analysis outcomes are added to these existing data objects, rather than stored in additional data objects. Output tables and plots are automatically exported to subdirectories, categorized by analysis type. In addition, SambaR contains tools which allow to subset (based on sample/locus names), subsample and intersect datasets (i.e. finding overlap between SNP datasets), and to detect small subsets of SNPs which are most informative with respect to population structure.

Here we describe SambaR and test the software on previously published SNP datasets. We also discuss new tools implemented in SambaR, including a) output plots which can help users to optimize their filtering settings, b) a Bayesian population assignment (BPA) test, c) the ‘distinctive clustering-score’, a metric for objective measurement of the distinctiveness of population clusters based on sample loadings on ordination axes, d) functions which can estimate genome wide heterozygosity from reduced representation sequencing libraries, and e) functions which extract and export reduced SNP panels of various sizes.

## Methods and materials

### Technical details

SambaR is implemented as an R package and can run on any operating system. The package has been tested on Windows, Linux and Mac computers. SambaR will install up to 2GB of dependencies (i.e. other R packages needed by SambaR for plotting and data analysis). Due to this dependency on other packages, for full use SambaR requires recent R versions (currently 4.0.0 or higher).

### SambaR pipeline

The SambaR pipeline consists of seven main functions (Fig. S1, Table S1):

- The ‘getpackages’ function installs and downloads dependencies. Optionally users can edit an automatically generated control file (‘mypackageslist.txt’) to prevent SambaR from attempting to install certain packages. The control file classifies packages into three categories: ‘essential’, ‘recommended’, and ‘optional’. Essential packages are required for SambaR to run without errors. Recommended packages are needed for key analyses.
- The ‘importdata’ function uses the read.PLINK function of the adegenet package (Jombart, 2008; Jombart & Ahmed, 2011) to import a SNP dataset (from binary PED/MAP format) into R and to store this data as a genlight object (Jombart, 2008) named ‘mygenlight’. Sample-specific and locus-specific information are stored in two auxiliary dataframes called ‘inds’ and ‘snps’ respectively (Fig. S1). The function will incorporate in the ‘inds’ dataframe sample information provided in an optional sample file. The function will also incorporate in the ‘snps’ dataframe read depth and positional information found in optionally provided vcftools and STACKS output files. Monomorphic sites present in the input data file will be excluded from subsequent analyses.
- The ‘filterdata’ function executes quality control. This function adds to the ‘inds’ and ‘snps’ dataframe boolean vectors (i.e. inds$filter, snps$filter and snps$filter2) which determine which samples and loci are included in subsequent analyses (Fig. S1). Current filter options include: missing data per locus, missing data per sample, minor allele count per locus, locus specific deviation from HWE, read depth per locus, read depth per sample, transitions vs transversions, and, if genome locations are provided by the user, spacing between SNPs. Relatedness between samples is estimated by the kinship coefficient (Waples et al, 2018) and by the KING-robust measure (Waples et al, 2018).
- The ‘findstructure’ function uses various R packages to perform principal components analysis (PCA) and principal coordinates analysis (PCoA), multi-dimensional scaling (MDS), discrimation analysis of principal components (DAPC, Jombart et al, 2010), correspondence analyses (CA), admixture analyses (using the R package LEA), and in addition generates structure-like plots with Bayesian population assignment (BPA) probabilities (see below). PCoA analyses are performed on three different types of genetic distance estimates: Nei’s genetic distance, Hamming’s genetic distance, and pairwise sequence dissimilarity. If the user provides samples locations (i.e. geographical coordinates), SambaR also generates geographical maps and in addition runs Tess3r (Caye et al, 2018) to perform tessellation analyses.
- The ‘calcdistance’ function generates population differentiation measures for all pairwise population comparisons. These estimates including D_xy_, fst_π_ (Hudson et al, 1992), Nei’s genetic distance and Weir & Cockerham 1984 Fst, the latter two generated using functions of the StAMPP package (Pembleton, 2013). In addition to genome wide estimates, the function also generates locus specific Fst estimates, using three different metrics (Nei, 1977; Weir & Cockerham, 1987; Wright, 1943, Supplementary methods). D statistics are generated using ABBA-BABA calculations described in Durand et al, (2011).
- The ‘calcdiversity’ function performs 1D and 2D site frequency spectrum (SFS) analyses, calculates nucleotide diversity and pairwise sequence dissimilarity estimates, and screens the genome for runs of homozygosity (using the R package detectRUNS, Filippo et al, 2018). If users provide the number of chromosomes to the nchroms flag (i.e. number of biggest scaffolds to include), SambaR will in addition generate karyotype plots (Gel & Serra, 2017) showing genome wide variation.
- The ‘selectionanalyses’ function uses the R packages Fsthet (Flanagan & Jones, 2018), OutFLANK (Whitlock & Lotterhos, 2014, 2015), PCadapt 4.1.0 (Luu et al, 2017, 2019) to search for SNPs under balancing or diversifying selection. The function also executes Fisher exact tests for associations between allele frequencies and population assignment.

### Plotting

During execution of SambaR’s main functions, results are automatically exported into ready-to-publish plots in four different file formats: eps, pdf, png, and, depending on the operating system, wmf. Layout settings, including font type, font size and colour coding matching population assignment, are coherent. Plots are generated with various settings allowing users to generate plots according to personal preferences. Function arguments allow users to easily customize colour coding and font type, as well as the size and location of the legend. Default font and dot sizes ensure readability even if plots are scaled down. Output files are stored in subdirectories named QC, Structure, Divergence, Diversity, Demography, Selection and Inputfiles. These subdirectories are located within a main directory called SambaR_output.

Several subsets of plots are automatically combined by SambaR into multi-tile figures and exported in the pdf format. For more advanced R users, SambaR provides a function to create custom multi-tile figures with user defined combinations of SambaR plots.

### List of R packages used by SambaR

Currently SambaR uses the following R packages to perform population-genetic analyses: adegenet (Jombart, 2008; Jombart & Ahmed, 2011), ape (Paradis & Schliep, 2018), detectRUNS (Filippo et al, 2018), FactoMineR (Lê et al, 2008), Factoextra (Kassambara & Mundt, 2019), HybridCheck (Ward & Van Oosterhout, 2016), LEA (Frichot & François, 2015), OutFLANK (Whitlock & Lotterhos, 2014, 2015), pcadapt (Luu et al, 2017, 2019), poppr (Kamvar et al, 2014), StAMPP (Pembleton et al, 2013), qvalue (Storey et al, 2019), tess3r (Caye et al, 2018), SNPRelate (Zheng et al, 2012), and zoo (Zeileis & Grothendieck, 2005).

For plotting, Sambar makes use of the R packages: circlize (Gu et al, 2014), colorspace (Zeileis et al, 2019), gplots (Warnes et al, 2019), grid (P. Murrell, 2005), gridGraphics (Paul Murrell & Wen, 2019), gridExtra (Auguie, 2017), karyoploteR (Gel & Serra, 2017), mapplots (Gerritsen, 2018), migest (Abel, 2019), plot3D (Soetaert, 2017), plyr (Wickham, 2011), RColorBrewer (Neuwirth, 2014), raster (Hijmans, 2019), rworldmap (South, 2011), scales (Wickham & Seidel, 2019), scatterplot3D (Ligges & Mächler, 2003), VennDiagram (Chen, 2018) and vioplot (Adler & Kelly, 2019).

### Analyses outside of the R environment which are supported by SambaR

SambaR also facilitates the usage of software outside of R. These include Admixture (Alexander et al, 2009), Bayesass (Mussmann et al, 2019), Bayescan (Foll & Gaggiotti, 2008), GCTA (Yang et al, 2011); PLINK (Chang et al, 2015; Gaunt et al, 2007; Purcell et al, 2007; for linkage disequilibrium, inbreeding and relatedness calculations) and Stairwayplot (Liu & Fu, 2015). SambaR does so by creating input files for these programs, such as site frequency spectrum vectors needed for Stairwayplot, and by generating plots from their output files.

### Highlighted features of SambaR

In the sections above we described the SambaR pipeline. In the following we will highlight particular features of SambaR, which include population-genetic tools uniquely implemented in SambaR.

#### Data filtering recommendations

As outlined above, the main purpose of SambaR’s ‘importdata’ function is to import SNP data into R. In addition, the ‘importdata’ function also generates plots which users can consult for choosing filter settings appropriate for their research questions.

Users are recommended to execute population structure analyses with a strict ‘snpmiss’ filter that sets the maximum proportion of missing data of retained SNPs close to zero. This prevents distortion of ordination plots due to variation in levels of missing data between samples (Fig. 1, S2–S3). The strictness of the SNP filter is however limited by the quality of the data, because a sufficient number of retained SNPs are needed to discern population structure. The ‘Data_quality’-plot (Fig. 1A) shows the number of retained SNPs as a function of missing data thresholds, and therefore allow users to choose the minimum threshold that is needed to retain the desired number of SNPs.

**Figure 1.**
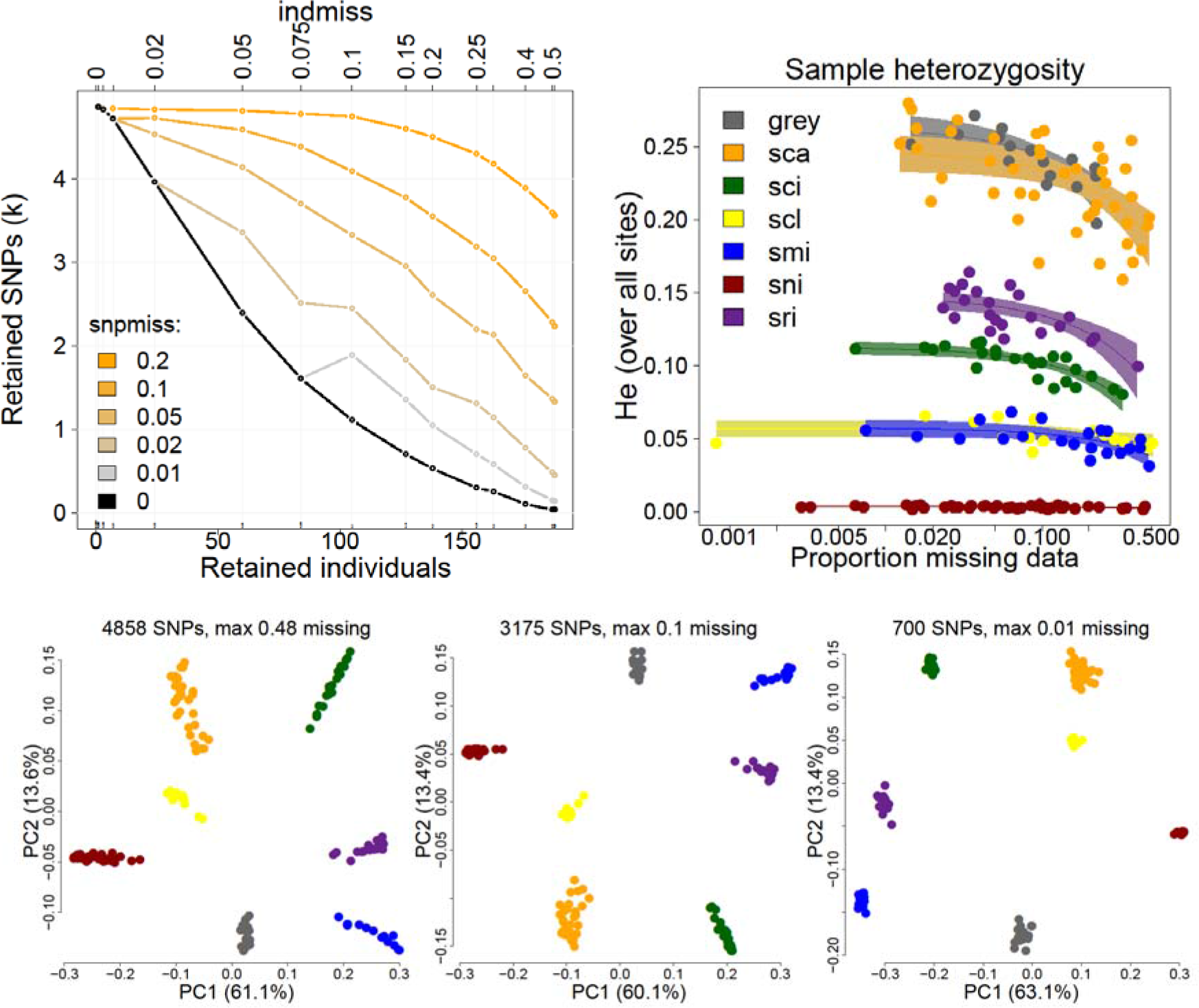
Filter settings and pcoa analyses. Analyses outcomes for a RADseq dataset of the island channel fox and the closely related mainland grey fox (Funk et al. 2017). **A**. Number of retained SNPs as a function of filter settings (snpmiss: maximum proportion of missing data points per SNP; indmiss: maximum proportion of missing data points per individual/sample). **B**. Sample heterozygosity vs proportion missing data. Inclusion of samples with high proportions of missing data leads to underestimates of genetic diversity. **C**. Principal coordinates analyses (PCoA) based on Hamming’s genetic distance using SNP datasets with different filter settings. Colour coding according to Funk et al (2017).

For genetic diversity analyses, SambaR users are recommended to exclude all samples for which the heterozygosity estimates are likely biased by relatively high proportions of missing data. These samples can be identified using the ‘Heterozygosity_vs_missingness’ plot (Fig. 1B).

SambaR performs population structure, diversity and differentiation analyses on a thinned dataset containing maximum one SNP per 500 bp (default settings). In contrast, selection analyses are performed on a non-thinned dataset, because the detection of linked outlier SNPs strengthens inference about selection. Although it is common practice to filter SNP datasets based on linkage disequilibrium considerations, SambaR users interested in genetic diversity and selection analyses are recommended to not thin their data set prior to importing the data into R, unless because of size limitations. Full, non-thinned, datasets enable estimation of genome wide heterozygosity (see below) and facilitate the generation of dense Manhattan plots.

#### Pairwise sequence dissimilarity, nucleotide diversity (π), Watterson’s theta and Dxy

The SambaR-function ‘calcpi’, which is invoked by several main functions, calculates for each pair of individuals pairwise sequence dissimilarity estimates (Supplementary Methods). These estimates are subsequently used to calculate several dependent population-genetic measures, including nucleotide diversity (π), Watterson”s theta, Tajima’s D, Fst_π_ and D_xy_ (Supplementary Methods).

#### Genome wide heterozygosity and π estimates inferred from RADseq datasets

SambaR generates estimates of genome wide heterozygosity (Fig. 2) to facilitate comparisons of genetic diversity between SNP datasets. Users can enable this estimation by providing input value to the nrsites flag of the ‘calcdiversity’. This input value should be an estimate of the total number of sequenced sites in the filtered sequencing read dataset from which the SNP dataset is derived. The calculation assumes that users did not select at maximum one SNP per read, filtered their genotype file prior to extracting of bi-allelic SNPs (not vice versa), and did not thin their data based on linkage disequilibrium calculations.

**Figure 2.**
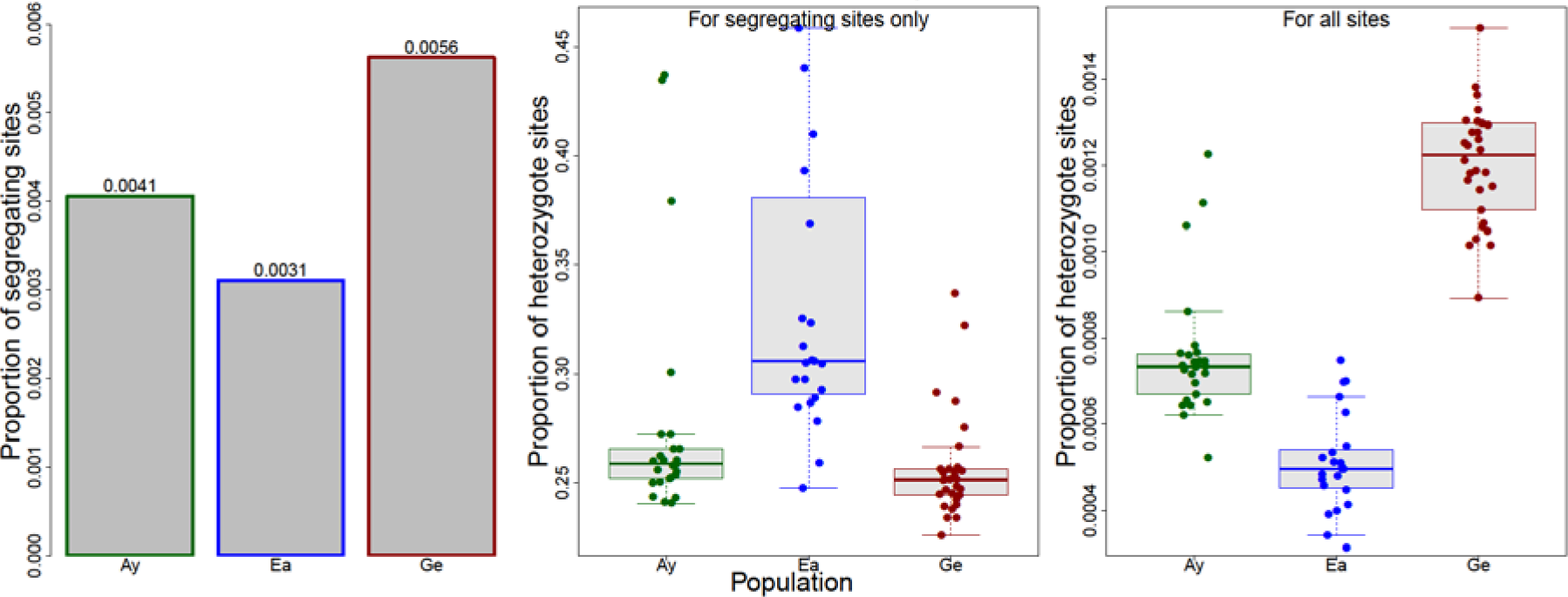
Genome wide heterozygosity estimates inferred from RADseq data. To facilitate comparability of genetic diversity estimates, SambaR generates estimates of genome wide heterozygosity from RADseq datasets. Shown are estimates of the proportions of segregating sites, sample heterozygosity over these segregated sites, and the final estimate of sample heterozygosity over all sequenced sites, for a dataset of three European roe deer populations (De Jong et al 2020). Colour coding according to De Jong et al (2020).

### DAPC analyses

Accurate execution of discriminant analyses of principle components (DAPC, Jombart et al, 2010), as implemented in the ‘adegenet’ package, depends on several parameters. These parameters include the number of principal components to retain, the number of clusters (K), and the inclusion or exclusion of a priori population structure information. SambaR explores DAPC parameter space by generating DAPC plots for various combinations of number of retained principal components, number of retained clusters (by default two to six), and inclusion and exclusion of a prior population structure information (Fig. 3). SambaR runs DAPC for five different values of retained principal components, one based on the a-score, and the other four corresponding to various percentages (i.e., 20, 50, 80 and 95%) of explained variance.

**Figure 3.**
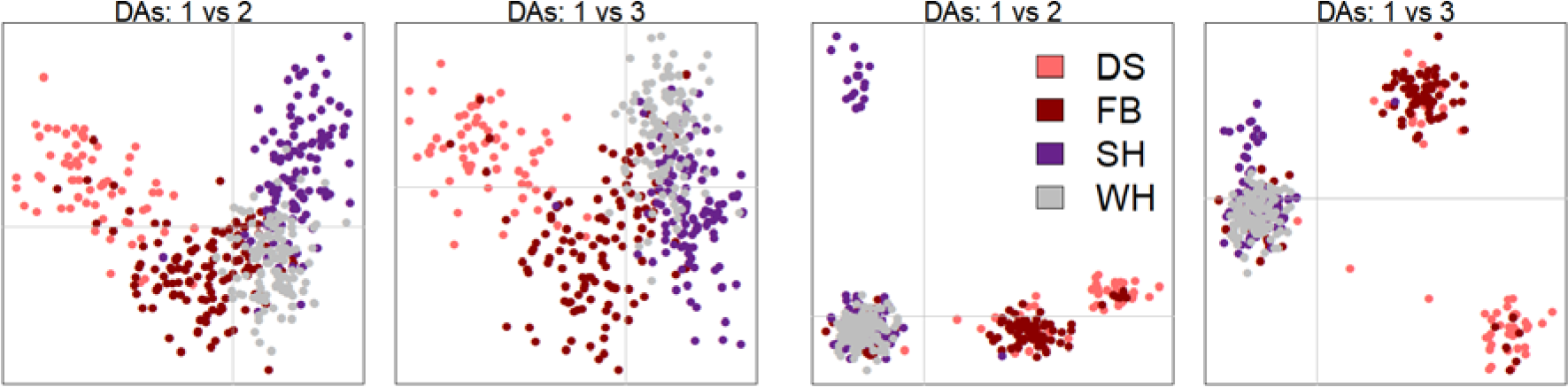
Goodness of fit between a priori defined and DAPC inferred population structure. DAPC-plots depicting ordination axes 1-3 for K=4 and for 33 retained principal components (left, 20 percent explained variance) and 238 retained principal components (right, 80 percent explained variance) for a dataset of 414 polar bears (Viengkone et al 2017). Limited overlap is observed between DAPC inferred clusters and a priori defined populations. Colour coding according to Viengkone et al (2017).

To guide users in selecting the most appropriate DAPC plot, SambaR generates the following summary statistics plots:

- a-score as a function of number of retained PCs, generated by the function optim.a.score() (Fig. S4)
- The estimated number of successful predictions as a function of number of retained PCs (i.e x-value for cross-validation, generated by the function xvalDapc(), Fig. S4)
- Explained variance as a function of number of retained PCs (Fig. S4)
- BIC-value as a function of a number of clusters (Fig. S4)
- Heatmaps depicting the overlap between predefined populations and DAPC inferred clusters (Fig. S5)

SambaR also performs a chi-squared test for the goodness of fit between a priori defined populations and DAPC inferred clusters for K equalling the number of a priori defined populations.

#### Selection of most informative SNPs

Depending on the study system, a relatively low number of highly informative SNPs is generally sufficient to assign individuals to populations (Von Thaden et al, 2020). These subsets of highly informative SNPs allow for low-cost determination of sample ancestry. Thus, there is a need for software to detect the most informative SNPs within a SNP dataset. SambaR contains a function, invoked by the ‘findstructure’ function, which detects SNPs with the highest standard deviation in population minor allele frequencies (given a predefined set of populations). Not included are SNPs for which data is lacking for one or more populations, nor SNPs of which the minor allele is missing in or more populations.

SambaR exports PED and MAP files of subsets of various sizes (50, 100, 150, and 250 SNPs) of these most informative SNPs, alongside estimates of population allele frequencies. SambaR also generates PCoA plots (Fig. 4) and Bayesian population assignment (BPA) test plots (Fig. S6) showing population structuring inferred from these reduced SNP panels.

**Figure 4.**
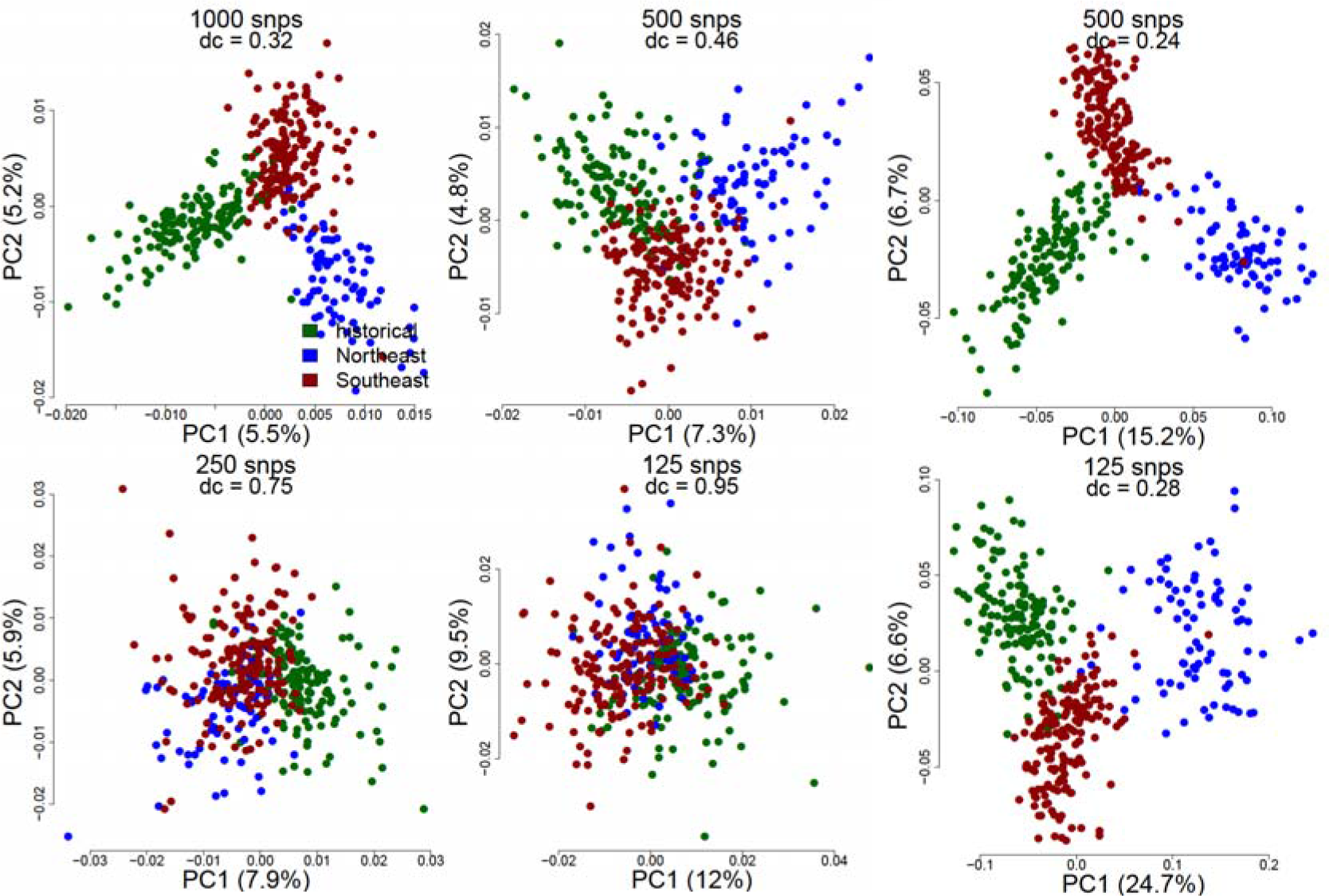
PCoA analyses on random SNP subsets (left/middle) and reduced SNP panels (right). PCoA analyses for SNP data subsets extracted from a ~22K SNP dataset of 394 coyotes (*Canis latrans*) sampled throughout the United States (Heppenheimer et al 2018). Left: PCoA plots for random subsets of 1000, 500, 250 and 125 SNPs. Right: PCoA plots for SNP subsets consisting of 500 and 125 SNPs with the highest standard deviation of minor allele frequency among populations. Plot titles indicate the dataset size and the dc-score, which measures the distinctiveness of population clusters (see main text for more information). Colour coding according to Heppenheimer et al (2018).

#### Bayesian population assignment test

SambaR contains a function, invoked by the ‘findstructure’ function, which calculates for each individual the posterior probability that this individual belongs to a set of predefined populations, using as input the population minor allele frequencies and individual genotypes. The question addressed by this Bayesian population assignment (BPA) test is: among a set of predefined populations, which is most likely to be the origin of a particular individual? The test is similar to previously published methods (Peatkau et al, 1995; Baudouin et al, 2004), with minor modifications (see Supplementary Methods for more details).

Because the BPA test assumes independency of loci, SambaR performs the calculations on a thinned dataset (if genomic locations of SNPs are provided). By default, the thinned dataset includes maximum 1 SNP per 500 bp. This threshold is arbitrary and can be changed by the user when running the ‘filterdata’ function.

Also excluded from the calculation are all SNPs for which one of either allele is missing in one or multiple populations, because these loci make the assignment probability converge to 0 or 1 immediately. A limitation of the BPA test is therefore that the test can only be applied to SNP datasets in which a sufficient number of SNPs have both alleles present in all populations. Depending on the study system, a few hundred biallelic SNPs might suffice (Fig. S6).

It has previously been reported that Bayesian posterior assignment probabilities are unreliable if population sample sizes are uneven (Peuchmaille, 2016). More generally, the reliability of the BPA test depends on the precision of the allele frequency estimates, which in turn depends on sample sizes. SambaR users are therefore advised to exercise caution when interpreting the BPA test results for datasets with a small or uneven number of individuals per population. Reliable estimation of population allele frequencies generally requires 30 or more individuals per population (Fung & Keenan, 2014).

Another potential shortcoming of the BPA test is circular reasoning. This occurs if the population specific allele frequency estimates are calculated based on a dataset which includes the individual for which the population assignment is being investigated. SambaR therefore recalculates population minor allele frequencies by excluding data for the investigated individual before running the BPA test (see Supplementary Methods).

#### dc-score

SambaR aims to facilitate an objective interpretation of ordination analyses by calculating the ‘dc-score’, which we introduce here. The dc-score, or ‘distinct clustering’-score, measures the overlap between population clusters in a two-dimensional space defined by the first and second ordination axis. The score is calculated by dividing the mean Euclidian distance of samples from their population centre by the mean Euclidian distances between population centres (Fig S7, Supplementary Methods). A dc-score close to zero indicates the absence of overlap between population clusters, whereas a dc-score greater than one indicates that the mean distance between population centres is smaller than the mean distance of samples to their population centre. Percentage of explained variance per ordination axis is not considered in the calculation.

## RESULTS AND DISCUSSION

### Data size limits and run time

To explore the data size limitations and run time of SambaR, we used SambaR to analyse online available whole genome sequencing (WGS) and reduced representation library (RRL) SNP datasets on three computers with different capacities. The findings indicate that the run time of SambaR, and whether it completes without encountering memory allocation errors, mainly depends on the capacities of the computer (Table S2). The run time estimates (Table S2) can provide guidance for users to match computational capacity to the dataset in question, or alternatively to filter down datasets to computational capacity.

On High Performance Clusters, SambaR can process datasets of more than one million SNPs and more than hundred individuals within hours. On ordinary desktop computers, datasets containing over 200K SNPs and over 100 individuals are likely to result in memory allocation errors, depending on the memory dimensions of the computer. Datasets of less than 100K SNP and less than 100 individuals are typically processed in less than an hour on an average desktop computer (Table S2).

### Data filtering recommendations

We explored the effect of data filter settings on the outcome of ordination analyses using a published RADseq SNP dataset of the island channel fox (*Urocyon littoralis*, Funk et al, 2016). PCoA analyses plots based on Hamming’s genetic distance using SNP datasets with a relaxed SNP filter threshold resulted in distorted ordination plots (Fig 1C), with sample loadings on ordination axes being a function of their proportion of missing data points (Fig S2). This distortion was not observed for a relatively small dataset of 700 SNPs, which were obtained after excluding all SNPs with maximum one percent missing data points. Similar findings were observed when running principal component analyses (PCA, Fig S3). These findings support SambaR’s recommendation to perform structure analyses with a relatively low number of high quality SNPs rather than with a high number of low quality SNPs.

For diversity analyses, in contrast, SambaR users are recommended to use a dataset which exhibits no relationship between the proportion of missing data points and heterozygosity per sample. For the island channel fox dataset, it can be argued that individuals with more than ten procent missing data should be excluded from the analyses (Fig S1B).

### Genome wide heterozygosity estimates inferred from RADseq datasets

To assess the accuracy of the genome wide heterozygosity estimation of SambaR, we compared SambaR’s estimate for two RADseq datasets against published WGS estimates.

Analysis of a European roe deer (*Capreolus capreolus*) RADseq ~50K SNP dataset (De Jong et al, 2020), derived from a total number of 8,196,980 sequenced sites (De Jong et al, 2020), returned genome wide heterozygosity estimates ranging between 0.09% and 0.14% per sample for a German roe deer population (Fig 2C). The WGS estimate of genome wide heterozygosity of a roe deer sample derived from the same locality equals 0.145% (De Jong et al, 2020).

Analysis of a Norwegian reindeer (*Rangifer tarandus*) RADseq ~80K SNP dataset (De Jong et al, in prep), derived from a total number of 9627819 sequenced sites, returned a genome wide heterozygosity estimate of ~0.19%, ranging between 0.14% and 0.21% among samples. The WGS estimate of genome wide heterozygosity of northern European reindeer equals 0.21% (Weldenegodguad et al, 2019).

More comparisons are needed to draw robust conclusions about the precision of SambaR’s estimates of genome wide heterozygosity, and users are encouraged to compare obtained estimates to estimates from the literature (if available).

### DAPC analyses

We calculated the goodness of fit between a priori defined populations and DAPC inferred populations for a RADseq dataset of 414 polar bears (Viengkone et al, 2017). DAPC analyses with 33 retained principal components, K = 4, and with prior population information, resulted in a graph similar to figure 1 in Viengkone et al (2017). DAPC analyses with 238 retained principal components, K=4, and without prior population information, resulted in a graph similar to figure 2 in Viengkone et al (2017). For both settings overlap between predefined populations and DAPC clusters was poor (Fig. S4), and this was reflected in the highly significant goodness of fit test p-values (X^2^ = 298, df = 9, p = 0 and X^2^ = 278, df = 9, p = 0).

Chi-squared tests for goodness of fit between DAPC inferred and three a priori defined European roe deer populations (De Jong et al, 2020) resulted in non-significant p-values (X^2^ = 4.35, df = 4, p = 0.36 for both 20% and 80% explained variance), indicating DAPC inferred clusters generally corresponded to the predefined population structure (Fig. S5).

### Selection of high informative SNPs and BPA test

We tested the power of reduced SNP panels to infer population structure in a dataset of 394 coyotes (*Canis latrans*) sampled throughout the United States (Heppenheimer et al, 2018). SambaR generates these reduced SNP panels by selecting the SNPs with the highest standard deviation in minor allele frequency across populations. PCoA analyses indicated that random subsets of ≤500 SNPs generally gave poor power in resolving population structure (Fig. 4). In contrast, PCoA analyses using small subsets of SNPs with the highest standard deviation in minor allele frequencies among populations, separated out predefined populations (Fig. 4). Similar findings were observed when comparing the results of BPA tests on random and non-random SNP subsets (Fig. S6). These findings illustrate that reduced SNP panels which are generated by selecting SNPs with high standard deviation in population minor allele frequencies, have the potential of low-cost population assignment.

### dc-score

To evaluate the usefulness of the dc-score, we compared the dc-score of the PCoA analyses on random and selected SNP subsets of the coyote dataset (Heppenheimer et al, 2018). A strong correlation was observed between dc-score and the size of the SNP dataset, ranging from 0.31 for 1000 SNPs to 0.95 for 125 SNPs (Fig. 4). For subsets comprised of most informative SNP’s, the dc-score was less dependent on number of retained SNPs, and ranged between 0.24 for 500 SNPs and 0.28 for 125 SNPs (Fig. 4).

## CONCLUSION

SambaR facilitates rapid population genetic analyses on biallelic SNP datasets by removing three major time sinks: file handling, software learning, and data plotting. In addition, SambaR provides a convenient platform for SNP data storage and management, guides users to adapt appropriate filter settings, and provides new tools. These newly developed utilities allow for generating reduced SNP panels, for generating structure-like plots with Bayesian populations assignment probabilities, for generating genome wide heterozygosity estimates from reduced representation libraries, and for objective interpretation of ordination analyses using the so called ‘distinct clustering’-score.

## Author’s contributions

MdJ and JdJ developed the software. MdJ wrote the paper and the manual, and developed the dc-score and the BPA-test. ARH and AJ provided funding, feedback/advice, and input to the writing.

## Acknowledgments

We thank Sofia Esteves da Silva, Erandi Bonillas Monge, Matt Newbould, Vania Fonseca da Silva, Daniel Moore, Sarah Mueller, Maria Nilsson-Janke, Thomas Parker, Dennis Schreiber, Yinhla Shihlomule, Biagio Violi, Magnus Wolf and Paige Yates for testing SambaR and providing feedback. We thank Tilman Schell and Christoph Sinai for installation instructions.

## Funding

This work was supported by the British Deer Society, by the Kenneth Whitehead Trust, by Hesse’s funding program LOEWE and by the Leibniz Association.

## Conflict of interest

none declared

## Data accessibility

Datasets used in this study, as well as the SambaR script, can be found at: https://github.com/mennodejong1986/SambaR

## SUPPLEMENTARY FILES

This file contains:

- Supplementary tables (Table S1–S2)
- Supplementary figures (Fig. S1–S7)
- Supplementary methods (1-7)

### SUPPLEMENTARY TABLES

**Table S1.**
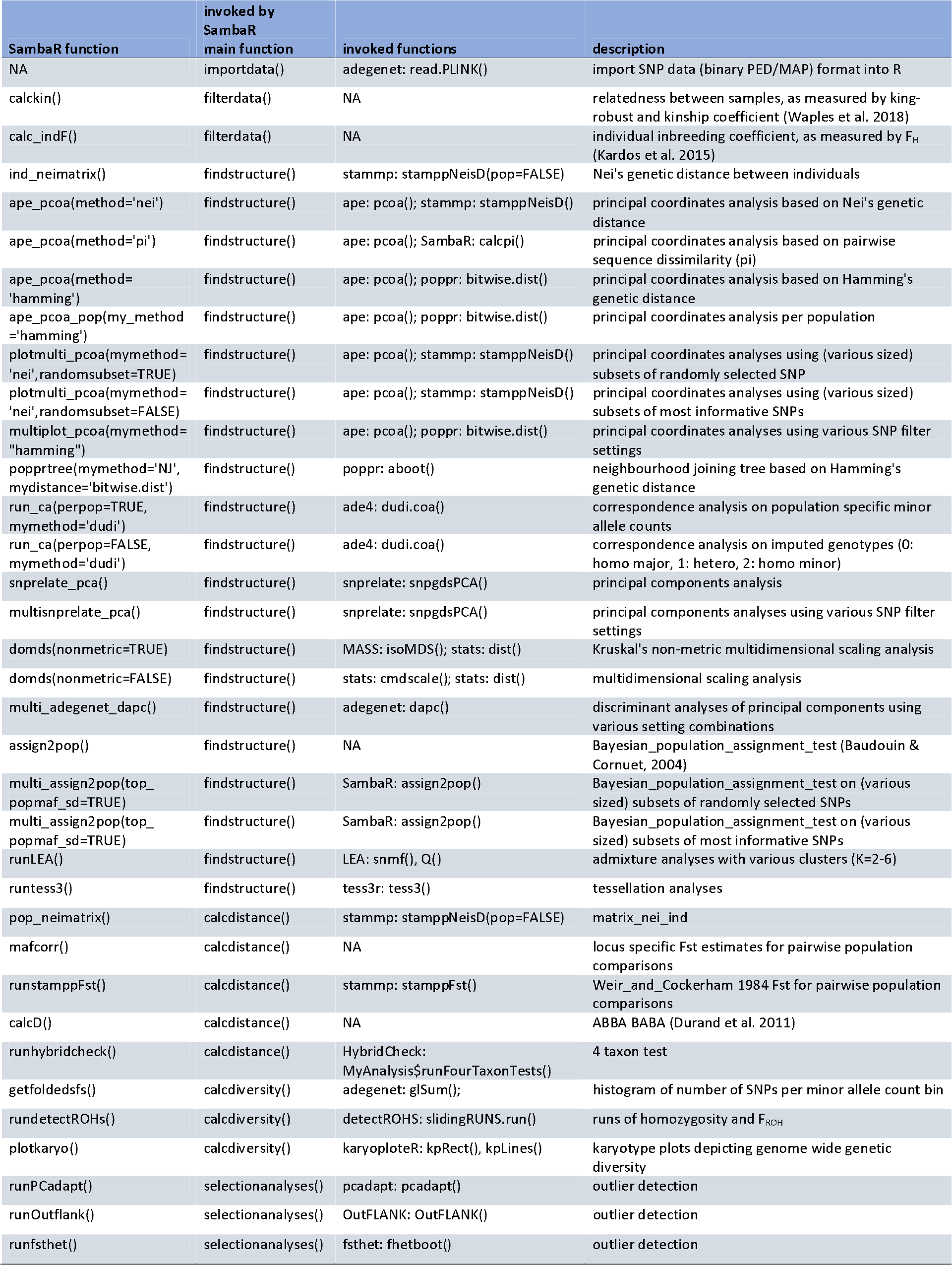
Overview of analyses included in the SambaR pipeline.

**Table S2.**
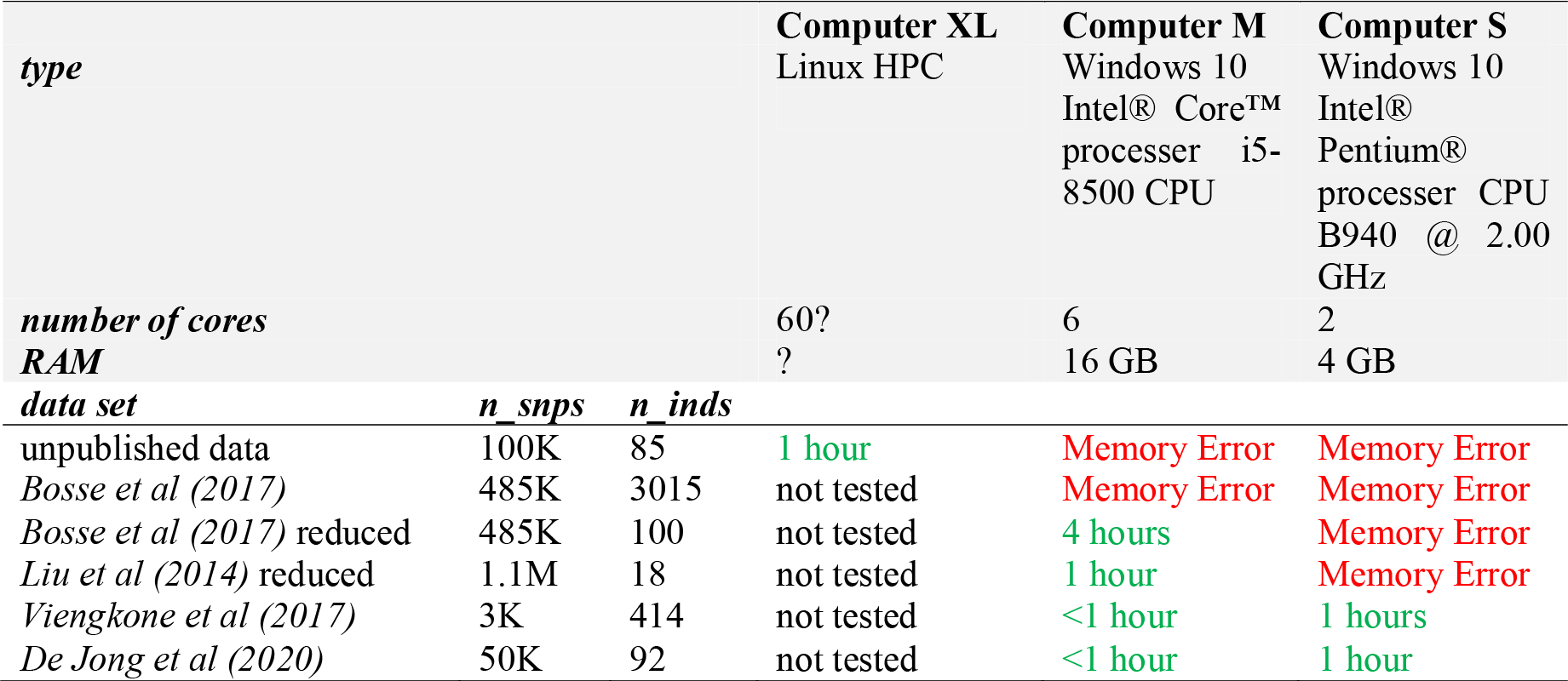
Run time of SambaR for datasets of various sizes (run time estimates do not include time needed for installation of dependencies)

### SUPPLEMENTARY FIGURES

**Figure S1.**
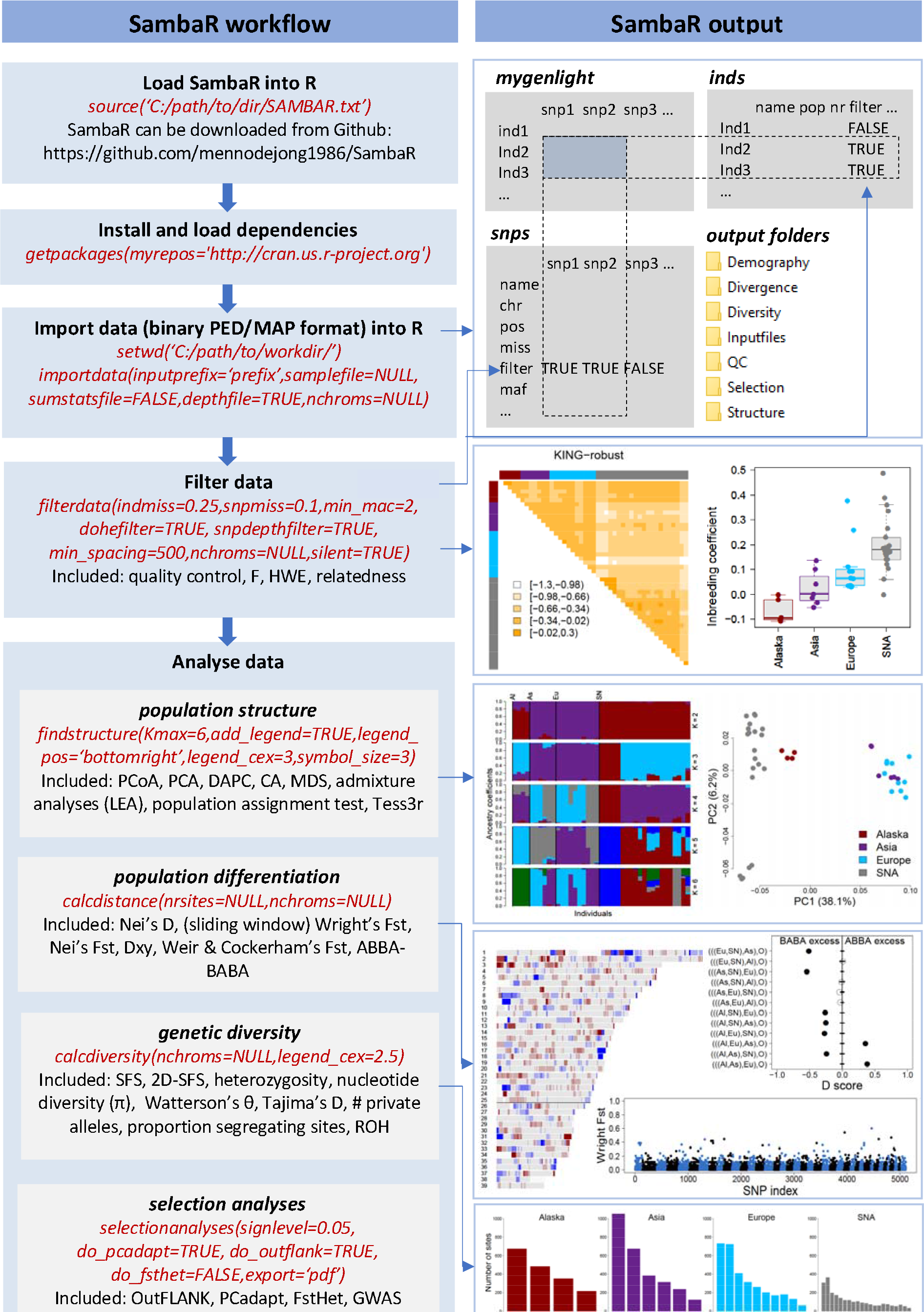
Schematic representation of the Sambar pipeline.

**Figure S2.**
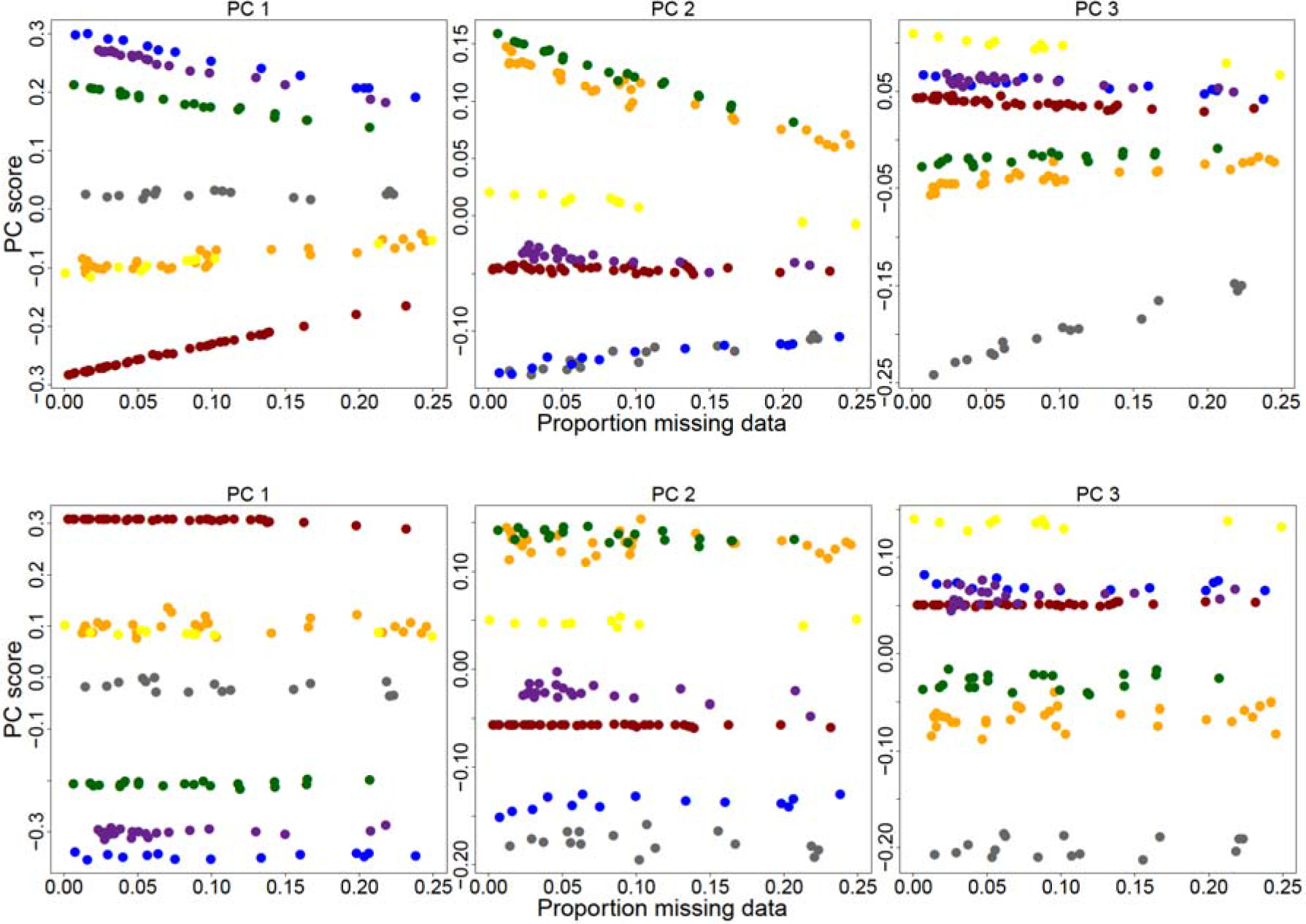
Sample loadings on PCoA axes vs proportion missing data. Sample loadings on PCoA ordination axes against proportion of missing data points per sample (measured against full SNP dataset) for a RADseq dataset of the island channel fox and the closely related mainland grey fox (Funk et al 2017). Above: sample loadings based on a dataset of 4,858 SNPs with maximum 50 percent datapoints per SNP. Sample loadings on the first three ordination axes are dependent of the quality of the sample. Below: sample loadings on a reduced dataset of 700 SNPs, obtained after removing SNPs with more than 1 percent missing datapoints. Sample loadings on the first three ordination axes are independent of the quality of the sample.

**Figure S3.**
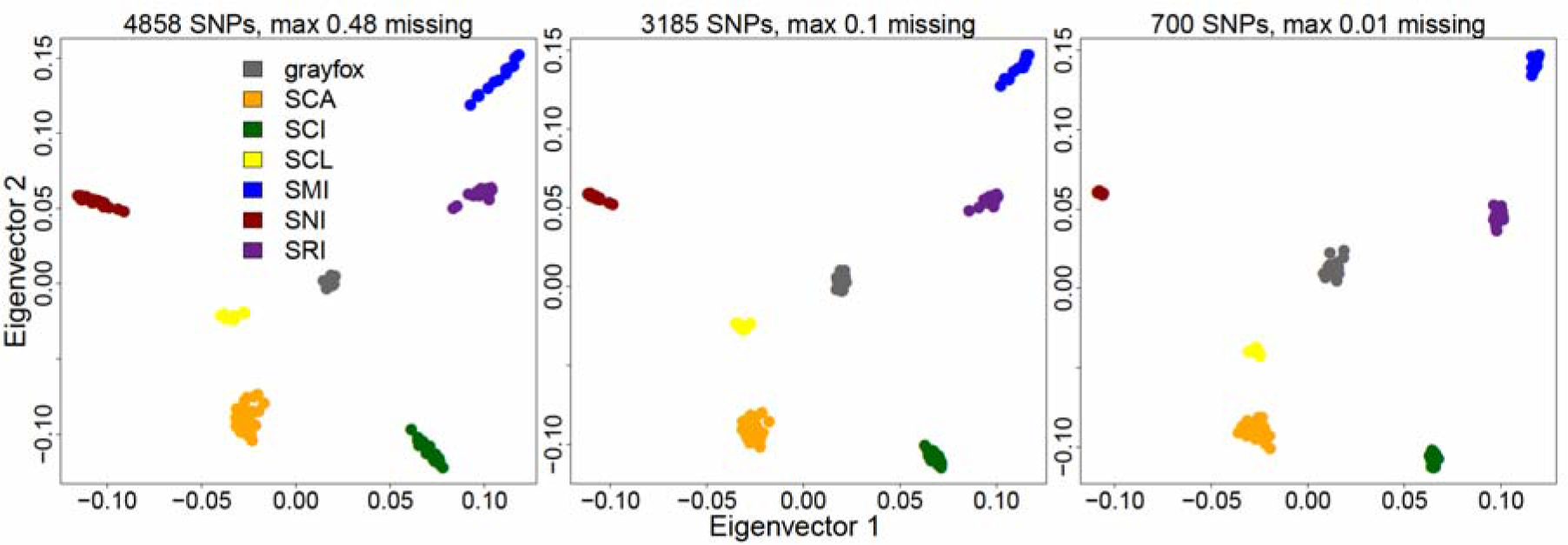
PCA analyses. PCA analyses, generated through functions implemented in the R package SNPrelate, for a RADseq dataset of the island channel fox and the closely related mainland grey fox (Funk et al 2017), based on various filter settings. As observed for PCoA analyses (figure 1), PCA analyses performed on datasets containing SNPs with high proportions (48% and 10%) of missing data points result in distorted ordination plots, which obscure the true distances between individuals.

**Figure S4.**
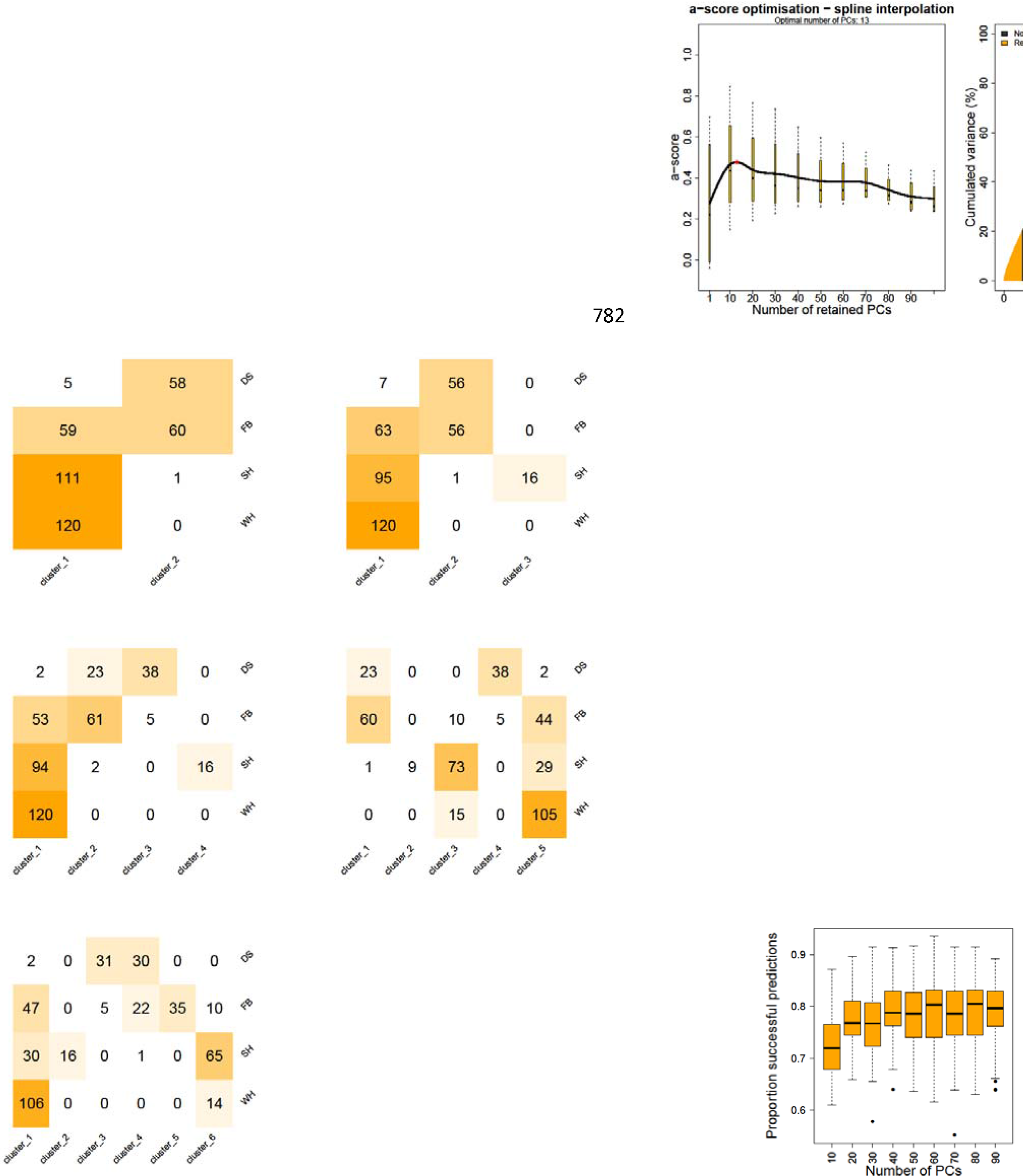
DAPC summary statistics generated by SambaR. The a-score, explained variance by principal components, BIC-score, heatmaps depicting overlap between a priori defined populations and DAPC inferred clusters for various K-values, and the x-value for cross validation, generated by the xvalDapc() function. Results are shown for analysis on a dataset of 414 polar bears (Viengkone et al. 2017), indicating limited overlap between a priori defined populations and DAPC inferred clusters.

**Figure S5.**
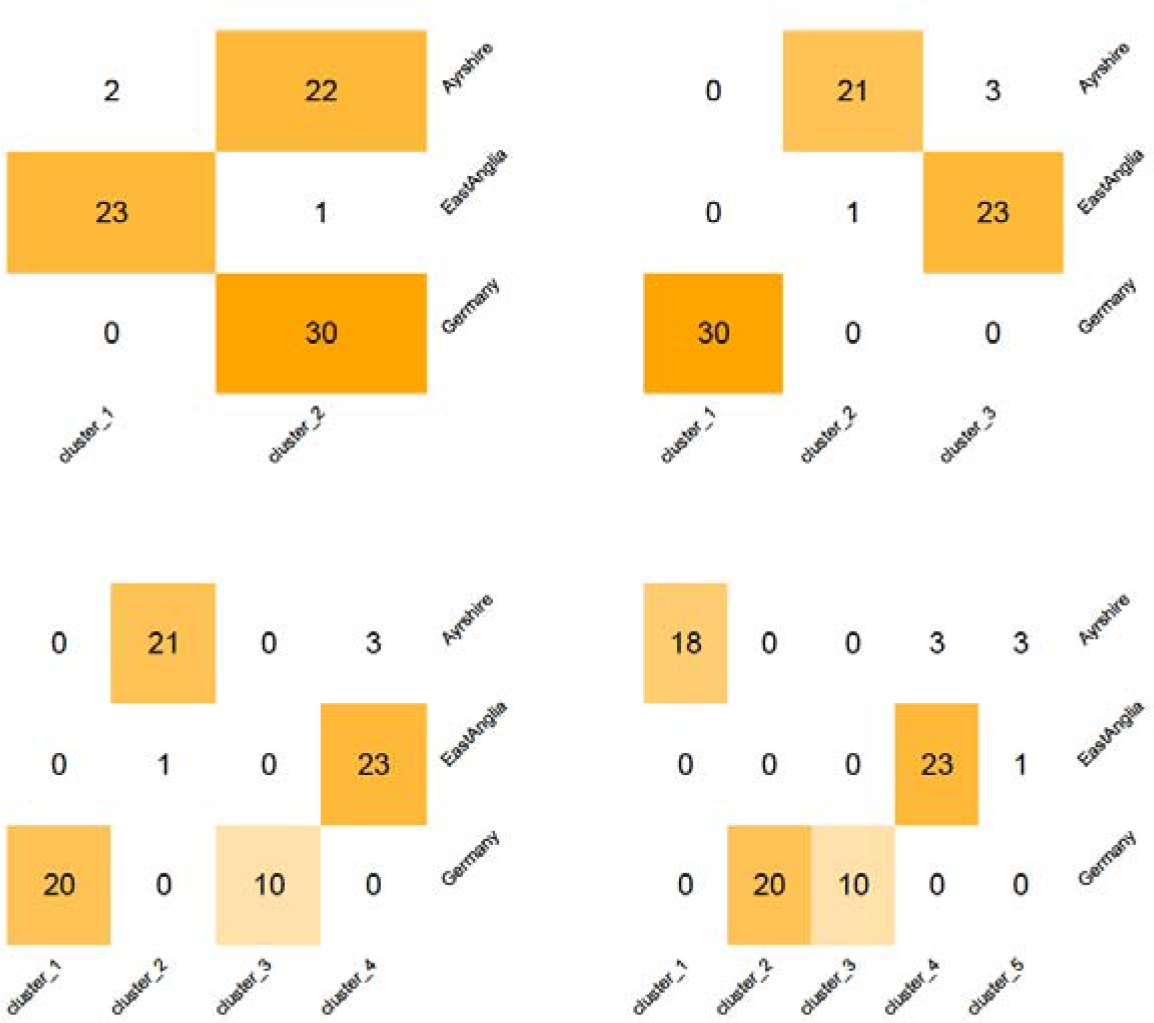
Correspondence between DAPC inferred clusters and predefined population structure. Heatmaps depicting overlap between a priori defined populations and DAPC inferred clusters for various K-values for a dataset of 94 roe deer (De Jong et al 2020). A chi-squared goodness of fit test between predefined populations and inferred DAPC clusters for K=3 (topright) returns a p-value of 0.36.

**Figure S6.**
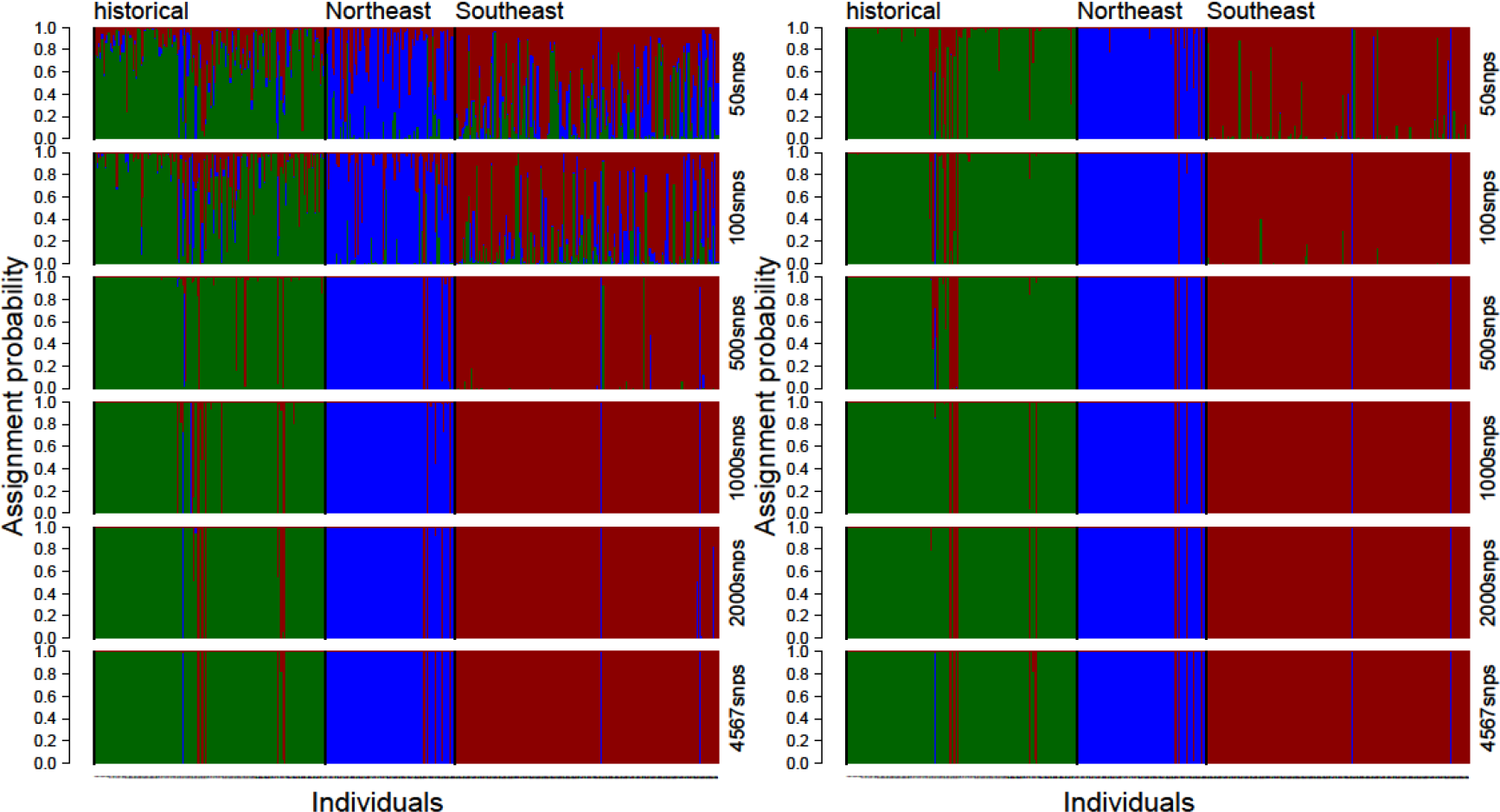
Bayesian population assignment (BPA) probabilities based on random SNP subsets (left) and based on subsets of SNPs selected by SambaR as most informative. Population assignment probabilities based on SNP data subsets extracted from a ~22K SNP dataset of 394 coyotes (*Canis latrans*) sampled throughout the United States (Heppenheimer et al 2018). Left: Assignment probabilities based on random subsets of 50, 100, 500, 1000, 2000 SNPs, as well as based on a full dataset of 4567 SNPs retained after quality control (indmiss=0.25, snpmiss=0.1). Right: Idem, but for subsets consisting of SNPs with the highest standard deviation of minor allele frequency among populations. Colour coding according to Heppenheimer et al. (2018).

**Figure S7.**
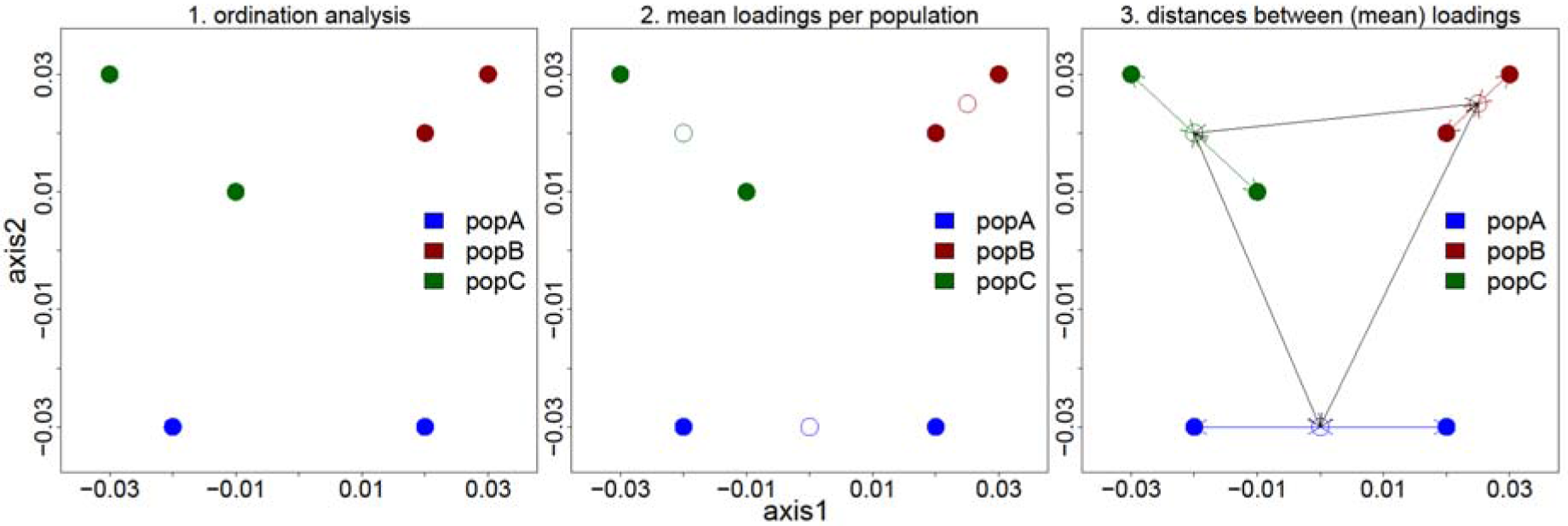
Visualization of the dc-score (‘distinct clustering’-score) calculation. 1. The dc-score is calculated based on the loadings of samples on the first two axes of an ordination analysis (e.g PCA or PCoA). 2. Population centres are defined as the mean loadings of populations on the ordination axes. 3. Euclidian distances between sample loadings and population centres (coloured lines), as well as Euclidian distances between population centres (black lines). The dc-score is defined as the mean length of the coloured lines divided by the mean length of the black lines.

### SUPPLEMENTARY METHODS

#### Supplementary methods 1: SambaR’s calculation of pairwise sequence dissimilarity, nucleotide diversity (π), Dxy, Watterson’s theta (Θ_W_) and Tajima’s D illustrated using an example dataset

Implemented in SambaR is the function ‘calcpi’, which is invoked by several of SambaR’s seven main functions. This function calculates observed pairwise sequence dissimilarities, followed by several population-genetic measures which depend on these estimates. Here we will describe the algorithm of the calcpi function using a small example dataset.

Consider the following genotype dataset for 3 individuals and 2 SNPs (in which 0 codes for homozygous major, 1 for heterozygous, 2 for homozygous minor, and NA for missing data):

|  |  | SNP1 | SNP2 |
| --- | --- | --- | --- |
| PopA | Ind1 | 0 | 1 |
| PopA | Ind2 | 2 | NA |
| PopB | Ind3 | 1 | 0 |

##### Pairwise sequence dissimilarity

SambaR calculates the number/proportion of differences and missing data points between and within individuals – taking into account that each sample pair provides four potential sequence comparisons (AB,Ab,aB,ab) – and subsequently estimates sequence dissimilarity, both for ‘between individual’-comparisons and ‘within individual’-comparisons:

|  | n_<br>comparisons | n_differences |  |  |  | n_missing_data_points |  |  |  | pairwise<br>sequence<br>O/(nsnps-M) |
| --- | --- | --- | --- | --- | --- | --- | --- | --- | --- | --- |
|  |  | SNP<br>1 | SNP<br>2 | total<br>l | mean<br>n | SNP<br>1 | SNP<br>2 | total<br>l | mean<br>n |  |
| Ind1 | 4 | 4 | NA | 4 | 1 | 0 | 4 | 4 | 1 | 1/(2-1)=1 |
| Ind1 | 4 | 2 | 2 | 4 | 1 | 0 | 0 | 0 | 0 | 1/(2-0)=0.5 |
| Ind2 | 4 | 2 | NA | 2 | 0.5 | 0 | 4 | 4 | 1 | 0.5/(2- |
| Ind1 | 1 | 0 | 1 | 1 | 1 | 0 | 0 | 0 | 0 | 1/(2-0)=0.5 |
| Ind2 | 1 | 0 | NA | 0 | 0 | 0 | 1 | 1 | 1 | 0/(2-1)=0 |
| Ind3 | 1 | 1 | 0 | 1 | 1 | 0 | 0 | 0 | 0 | 1/(2-0)=0.5 |
| total | 15 |  |  | 12 |  |  |  | 9 |  |  |

##### Nucleotide diversity (π)

Nucleotide diversity (π), which is the expected proportion of differences between any two randomly drawn sequences, is calculated as:

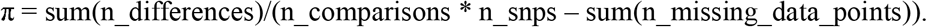

For the overall example dataset, π equals: *12/(15*2 – 9) = 12/21 = 0.5714*.

SambaR also estimates *π* per population:

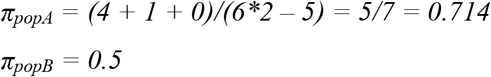

##### Dxy

SambaR obtains an estimate of *Dxy*, the mean proportion of differences between two sequences randomly drawn for two different populations, by calculating the mean number of pairwise sequence dissimilarity over all possible sample pairs.

For the example dataset, Dxy for pop1 and pop2 is obtained from averaging the pairwise sequence dissimilarity estimates obtained for Ind1-Ind3 and Ind2-Ind3:

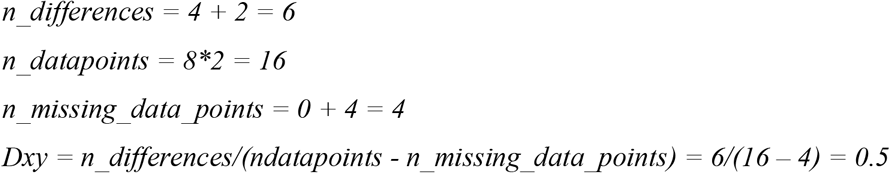

##### Fst_π_

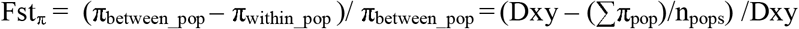

Fst_π_ for population pair popA and popB in the example dataset equals (see also the sections on nucleotide diversity and Dxy):

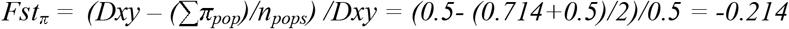

Negative Fst_π_ values indicate that individuals within populations are, in terms of their genotypes, more different from each other than individuals from different populations.

##### Watterson’s theta (θ_W_)

SambaR calculates Watterson’s estimate of theta (Θ_W_) by dividing the number of segregating sites within a population by the harmonic number (an). The number of segregating sites (S) is defined as the number of SNPs with a minor allele frequency above 0. The average number of sequences (n), needed to calculate the harmonic number (a_n_), is estimated as twice the number average number of individuals with non-missing data per SNP.

For the example dataset, Watterson’s theta for all individuals combined equals:

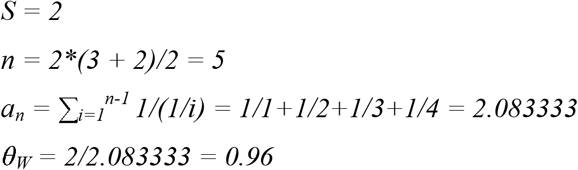

Hence for each pair of sequences (consisting of 2 SNPs) 0.96 sites are expected to differ (assuming neutrality). SambaR reports estimates per site (θ_W_persite_), which equals 0.48 for the numerical example.

##### Tajima’s D

SambaR calculates Tajima’s D scaled per single site as:

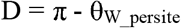

For the example dataset, SambaR’s Tajima’s D estimate (for all individuals combined) equals: *0.5714 – 0.48 = 0.0914*

SambaR also calculates significance using a chi squared test on observed and expected number of differences. For the example dataset (all individuals combined) the calculation becomes:

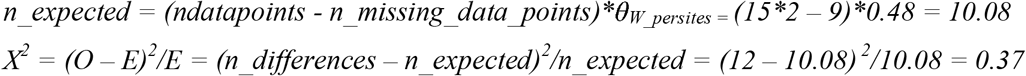

Next SambaR uses the R base function pchisq(df=1,lower.tail=FALSE) to derive the corresponding p-value:

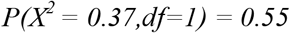

### Supplementary methods 2: SambaR’s estimation of genome wide heterozygosity

If users provide to the nrsites flag of SambaR’s calcdiversity function (default is NULL), an estimate of the total number of sequenced sites from which the SNP data is derived, SambaR will estimate genome wide estimates (e.g. genome wide heterozygosity and genome wide nucleotide diversity).

For RADseq data, the number of sequenced sites, here denoted as ‘N_total’, equals the combined length of all polymorphic as well as monomorphic stacks which passed filter settings, and can be obtained from the sumstats summary file.

For resequencing data, N_total equals the total number of monomorphic and polymorphic sites of the filtered data set. This number can for example be obtained by counting the number of lines in the VCF file, excluding the header section. The precondition is that the VCF file contains both monomorphic and polymorphic sites. If the VCF file is generated with bcftools, users should not include the -v option when running the bcftools calls command, otherwise monomorphic sites will not be outputted.

SambaR estimates genome wide heterozygosity in three steps:

1. SambaR determines ‘N_seg’, the number of segregating sites within the population to which the individual belongs.
2. SambaR calculates ‘He_seg’, the proportion of heterozygous sites within an individual for those segregating sites.
3. SambaR calculates genome wide heterozygosity using the formula: genome_He = He_seg * N_seg/N_total

### Supplementary methods 3: SambaR’s estimation of locus specific Fst-values, illustrated using an example dataset

Differences in minor allele frequencies (MAF, or p) between populations can be summarized using a Fst-metric. SambaR relies upon the stamppFst function (of the package ‘Stampp’) to generate Weir & Cockerham 1984 Fst estimates, along with associated significance values. This function generates for each population pairwise comparison one estimate for all SNPs combined.

In addition, SambaR uses built-in functions to calculate Fst estimates for each single SNP. SambaR generates three different Fst measures:

- Fst = Var(p)/(p(1-p)) Wright, 1943
- Fst = (H_T_-H_S_)/H_T_ Nei, 1977
- Fst = (f0 – f1)/(1 – f1) Cockerham & Weir, 1987

For illustrative purposes we will consider again the following genotype dataset for 3 individuals and 2 SNPs (in which 0 codes for homozygous major, 1 for heterozygous, 2 for homozygous minor, and NA for missing data):

|  |  | *SNP1* | *SNP2* |
| --- | --- | --- | --- |
| *PopA* | *Ind1* | 0 | 1 |
| *PopA* | *Ind2* | 2 | NA |
| *PopB* | *Ind3* | 1 | 0 |

#### Wright 1943 Fst

This measure was first published by Sewall Wright in 1943 (Wright, 1943, see equation 48 and the following lines).

*Var(p)*, also denoted as s_p_^2^, or σ_p_, denotes the variance in minor allele frequencies among populations. If p denotes the minor allele frequency, and if subscripts 1 and 2 denote either of the two populations considered in a pairwise population comparison, this estimate can be described as:

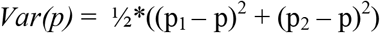

To standardize this estimate (i.e. make it comparable among SNPs regardless of their mean minor allele frequencies), the value it divided by the maximum value. It can be mathematically shown that for population pairs which are fixated for different alleles (i.e. p_1_=1 and p_2_=0 or vice versa), *Var(p)* equals: p(1-p). Therefore a standardized estimate is given by:

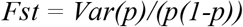

For SNP2 in the example dataset above, the Var(p) and Fst estimates equal:

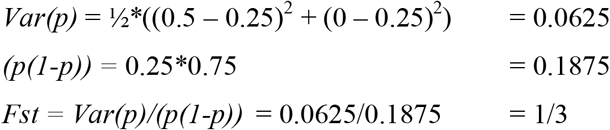

#### Nei 1977 Fst

This measure was developed by Nei (1977) for multi-allelic loci.

*H_T_* denotes the expected heterozygosity in the overall dataset (total population).

*H_S_* denotes the expected heterozygosity in the subpopulations.

If p denotes the minor allele frequency, and if subscripts 1 and 2 denote either of the two populations considered in a pairwise population comparison, these estimates are equal to:

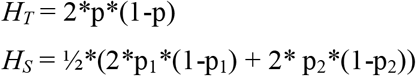

For SNP2 in the example dataset above, the H_T_ and H_S_ estimates are equal to:

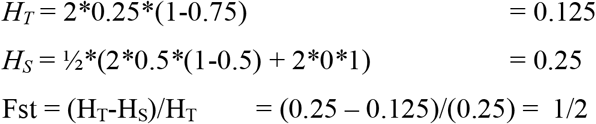

#### Cockerham & Weir 1987 Fst

This measure was published by Cockerham and Weir in 1987, and is not be confused with their Fst metric published in 1984. The original notation used by Cockerham and Weir (1987) was: Fst = (θ_1_ – θ_2_)/(1 – θ_2_). Here we denote θ_1_ as f_0_ and θ_2_ as f_1_.

*f0* denotes the probability that two randomly drawn alleles from within a (sub)population are identical.

*f1* denotes the probability that two randomly drawn alleles from the overall population (the total or metapopulation) are identical.

If p denotes the minor allele frequency, and if subscripts 1 and 2 denote either of the two populations considered in a pairwise population comparison, these estimates are equal to:

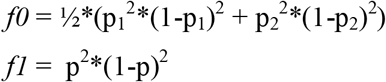

For SNP2 in the example dataset above, the f0 and f1 estimates are equal to:

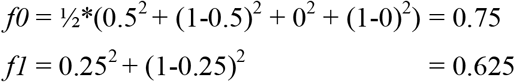

Hence, there is a 75% probability of drawing two identical alleles when pooling from within one of the populations, compared to a 62.5% probability when drawing two alleles from the metapopulation.

The Fst estimates for SNP2 becomes:

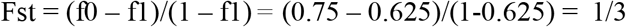

Note that from its definition it follows that *f1* is an estimate of expected homozygosity in the populations in the absence of population structure. Expected heterozygosity is therefore simply:

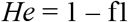

The ‘selectionanalyses’-function of SambaR will generate Fdist plots, showing locus specific Fst estimates on the y-axis against He (= 1 – f1) estimates on the x-axis.

### Supplementary methods 4: SambaR’s estimation of inbreeding coefficient F

SambaR uses two methods to calculate individual inbreeding coefficient F, the probability that the two alleles at any locus of a diploid individual are identical by descent (IBD): F_H_ (Kardos et al., 2015) and the here defined Fπ (see below).

#### F_H_

F can be defined as the proportion of expected heterozygous sites (in case of no inbreeding) which are homozygous due to inbreeding. In formula:

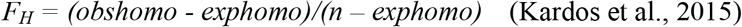

In which:

n: total number of sites (for an individual), excluding missing data points
obshomo: observed number of homozygous sites (for an individual)
exphomo: expected number of homozygous sites (for an individual), estimates as:

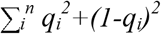

in which q_i_ denotes the minor allele frequency in the population to which the individual belongs, which is derived from all individuals in the population except for the individual for which F is calculated.

SambaR evaluates significance by applying a chi-squared test on observed and expected number of homozygous sites.

Consider, as an example, a diploid individual A which is genotyped at 10 sites, of which 6 are heterozygous. Through selfing individual A produces individual B. The most likely genotype of individual B consists of (4 + 0.5*6 =) 7 homozygous sites.

If all other individuals in the population are not inbred, and on average have the same number of homozygous and heterozygous sites as individual A, then the expected number of homozygous sites equals 4. The inbreeding coefficient of individual B therefore equals:

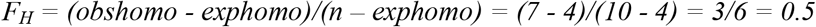

The *F_H_* value of 0.5 correctly indicates that individual B results from selfing. It also correctly indicates the proportion of autozygous sites of individual B (i.e. proportion IBD). Assuming the homozygous sites of individual A are IBS (identical by state) but not identical by descent (IBS), the most likely number of autozygous sites of individual B equals (0.5*10 =) 5.

Significance is evaluated using a chi-squared test:

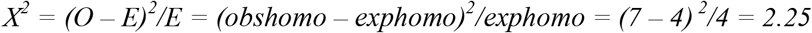

SambaR uses the R base function pchisq(df=1,lower.tail=FALSE) to derive the corresponding p-value:

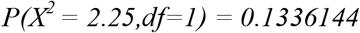

#### Fπ

The second method SambaR uses to estimate heterozygosity is based on a comparison between the heterozygosity (He) of an individual (j) and the mean pairwise sequence dissimilarity (here defined as πj) of this individual with all other individuals in the population:

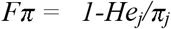

SambaR evaluates significance by applying a chi-squared test on observed and expected number of heterozygous sites, in which the number of expected heterozygous sites is defined estimated by πj times the number of sites.

Consider, as an example, an individual with a heterozygosity of 0.23 and a mean pairwise dissimilarity with other individuals of the population of 0.25. SambaR estimates the inbreeding coefficient for this individual as:

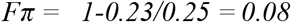

If the heterozygosity and nucleotide diversity estimates are based on a dataset of 1000 sites, a chi-squared test returns:

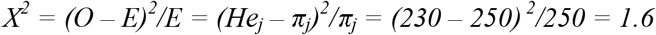

SambaR uses the R base function pchisq(df=1,lower.tail=FALSE) to derive the corresponding p-value:

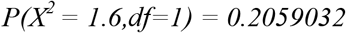

### Supplementary methods 5: SambaR’s Bayesian population assignment (BPA) test, illustrated using an example dataset

The BPA test addresses the question: among a set of predefined populations, which population is most likely to be the origin of a particular individual? The BPA test calculates the probability that an individual originates from each of the predefined populations, given the observed genotype of the individual and given the observed population allele frequencies.

Consider, as an analogy, two jars. Jar A contains 10 red and 90 white balls. Jar B contains 1 red and 99 white balls. A ball is drawn from one of these jar, and the ball is red. From which bottle has the ball been drawn? According to Bayes Rule, the posterior probabilities are:

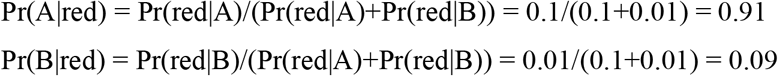

Hence, there is a 91% probability the ball has been drawn from jar A, and a 9% probability the ball has been drawn from jar B.

The BPA test uses a similar algorithm to calculate the probabilities that a sample has been drawn from each of the set of predefined populations:

Let Hkj denote the probability that individual k belongs to population j, let MAFij denote the minor allele frequency of locus i in population j (excluding the genotype of individual k), let Oi denote the genotype of individual k for locus i (i.e. number of minor alleles (0, 1, or 2)), and let k denote the number of predefined populations. For any given locus, the probability that an individual belongs to population j can be described as:

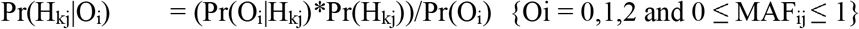

in which:

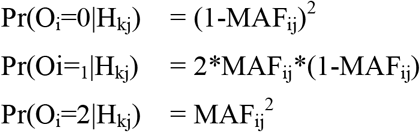

If assuming a flat prior distribution (i.e. Pr(H_kj_) = 1/j), the formula for Pr(H_kj_|O_i_) can be simplified to:

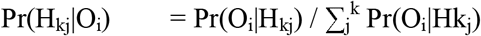

To calculate the probability for the nth locus, SambaR uses a recursive formula:

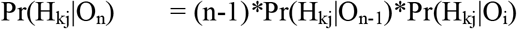

As an example, consider the following genotype dataset for 10 individuals and 2 SNPs (in which 0 codes for homozygous major, 1 for heterozygous, 2 for homozygous minor, and NA for missing data):

|  |  | *SNP1* | *SNP2* |
| --- | --- | --- | --- |
| *PopA* | *Ind1* | 2 | 2 |
| *PopA* | *Ind2* | 0 | 1 |
| *PopA* | *Ind3* | 1 | 0 |
| *PopA* | *Ind4* | 0 | 0 |
| *PopA* | *Ind5* | 0 | 2 |
| *PopB* | *Ind6* | 1 | 2 |
| *PopB* | *Ind7* | 2 | 2 |
| *PopB* | *Ind8* | 2 | 2 |
| *PopB* | *Ind9* | 1 | 1 |
| *PopB* | *Ind10* | 2 | 0 |

All individuals have been assigned to a set of two populations, PopA or PopB. Ind1 has been assigned to PopA, but what is the probability it does indeed belong to PopA (assuming the other population assignments are correct)?

The population minor allele frequencies, not considering Ind1, are:

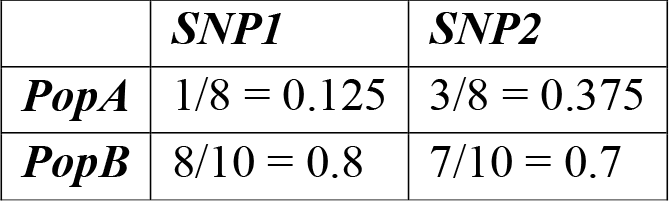

|  | *SNP1* | *SNP2* |
| --- | --- | --- |
| *PopA* | 1/8 = 0.125 | 3/8 = 0.375 |
| *PopB* | 8/10 = 0.8 | 7/10 = 0.7 |

For both SNPs, Ind1 is homozygous for the minor allele (i.e. genotype 0).

Let Pr_sum denote: (Pr(2|popA)+Pr(2|popB).

The posterior probabilities for SNP1 and SNP2 are equal to:

SNP1:

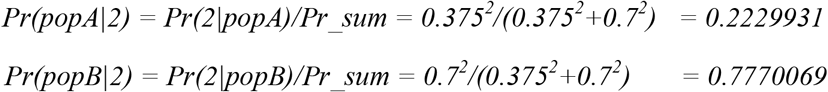

SNP2:

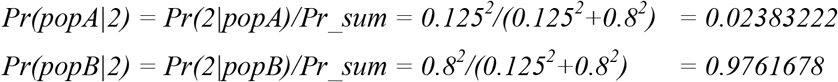

The combined posterior probabilities are equal to:

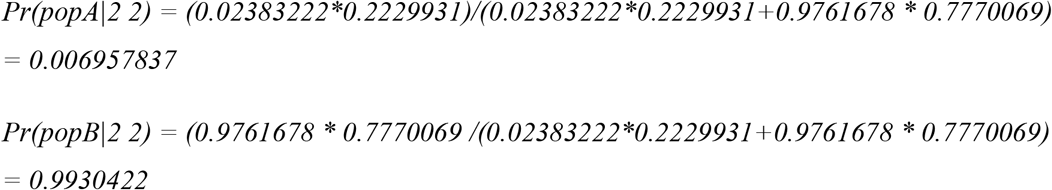

In conclusion, given the observed genotype of Ind1 and the observed allele frequencies in PopA and PopB, Ind1 belongs with 99.3% probability to PopB and with 0.7% probability to PopA, suggesting that the original population assignment is incorrect.

### Supplementary methods 6: The ‘distinct clustering’-score (dc-score) algorithm, illustrated using an example dataset

SambaR aims to facilitate objective interpretation of ordination analyses by calculating a ‘dc-score’ for PCA, PCoA, CA and DAPC analyses. The dc-score, or ‘distinct clustering’-score, measures the overlap between population clusters in a 2-dimensional space defined by the first and second axis of an ordination analysis (e.g. PCA and PCoA).

The algorithm behind the dc-score is as follows:

Let p1i_j and p2i_j denote the loading of sample i, belonging to population j, on the first and second ordination axes, and let nj denote the number of individuals belonging to population j.

The mean loadings (population centres) for population j are defined as:

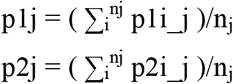

The mean distance dj of samples belonging to population j from the population centre of population j is defined as:

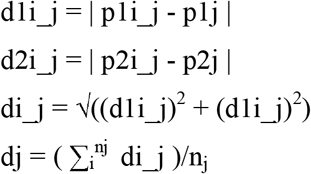

The mean distance of all samples from their population centres, given n_z_ populations, is defined as:

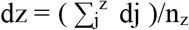

The distance between population centres for population pair k, dk, is defined as:

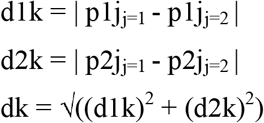

The mean distance between population centres for all n_m_ populations pairs, dm, is defined as:

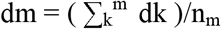

The dc-score is defined as:

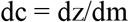

As an example, consider an ordination analysis on six samples (from three populations) returning the following output (Fig. S7):

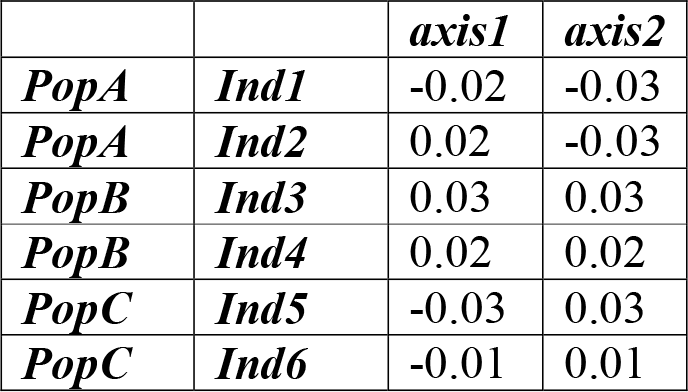

To calculate the dc-score, SambaR derives the mean loadings per population (‘population centres’):

To calculate the dc-score, SambaR derives the mean loadings per population ('population centres'):
|  | *axis1* | *axis2* |
| --- | --- | --- |
| **PopA** | 0 | -0.03 |
| **PopB** | 0.025 | 0.025 |
| **PopC** | -0.02 | 0.02 |

Thirdly, SambaR calculates the mean distance between sample loadings and population centres, and between population centres, and calculates the dc-score as the ratio between both distances:

|  |  | *axis1* | *axis2* | *distance* |  | *distance* | *dc (dz/dm)* |
| --- | --- | --- | --- | --- | --- | --- | --- |
| **PopA** | **Ind1** | -0.02 | -0.03 | 0.02 | **popA-** | 0.06041523 |  |
| **PopA** | **Ind2** | 0.02 | -0.03 | 0.02 | **popA-** | 0.05385165 |  |
| **PopB** | **Ind3** | 0.03 | 0.03 | 0.007071068 | **popB-** | 0.04527693 |  |
| **PopB** | **Ind4** | 0.02 | 0.02 | 0.007071068 |  |  |  |
| **PopC** | **Ind5** | -0.03 | 0.03 | 0.014142136 |  |  |  |
| **PopC** | **Ind6** | -0.01 | 0.01 | 0.014142136 |  |  |  |
| **mean** |  |  |  | dz = |  | dm = | 0.258319 |

### Supplementary methods 7: SambaR’s calculation of the site frequency spectrum (SFS), illustrated using an example dataset

SambaR generates site frequency spectrum (SFS) vectors in two steps:

- The first step involves counting for each SNP and for each population the number of minor allele copies. This counting is performed using the glSum function of the adegenet package. The minor allele is defined based on the total dataset. As a consequence, some populations can carry SNPs with more minor allele copies than major allele copies. If so, SambaR subtracts for these particular SNP the number of minor allele copies from the total number of observed allele copies (ignoring missing data points). This latter number is computed with the glNA-function of the adegenet package.
- The second step consists of binning and counting SNPs based on their number of minor allele copies.

By default, Sambar includes all SNPs in the calculation, also SNPs which did not pass filter settings. In contrast, it does not include individuals which did not pass the filter settings. The SFS vector generated by SambaR does not include sites with zero copies of the minor allele. As a result, the length of the SFS-vector for each population equals the number of retained individuals within this population.

Consider, as an example, the following genotype dataset for 7 individuals and 4 SNPs (in which 0 codes for homozygous major, 1 for heterozygous, 2 for homozygous minor, and NA for missing data):

|  |  | *SNP1* | *SNP2* | *SNP3* | *SNP4* |
| --- | --- | --- | --- | --- | --- |
| *PopA* | *Ind1* | 1 | 1 | 0 | NA |
| *PopA* | *Ind2* | 2 | 0 | 0 | 0 |
| *PopA* | *Ind3* | 2 | 0 | 0 | 2 |
| *PopA* | *Ind4* | 0 | 0 | 0 | 0 |
| *PopB* | *Ind5* | 1 | 1 | 2 | 0 |
| *PopB* | *Ind6* | 0 | 2 | 2 | 1 |
| *PopB* | *Ind7* | 0 | 0 | 2 | 0 |

#### Step 1. Count number of minor allele copies

Counting the number of minor allele copies results in the following data:

**Step 1. Count number of minor allele copies** Counting the number of minor allele copies results in the following data:
|  | *SNP1* | *SNP2* | *SNP3* | *SNP4* |
| --- | --- | --- | --- | --- |
| *PopA* | 8-5=3* | 1 | 0 | 2 |
| *PopB* | 1 | 3 | 6-6=0* | 1 |

\*Due to population structure, it can occur that within a population the minor allele (which is defined relative to the metapopulation) has in fact more copies than the minor allele. If that is the case, for this population and for this particular SNP the number of major alleles copies are counted (rather than number of minor allele copies).

#### Step 2. Binning SNPs based on their number of minor allele copies

The second step is to bin and count the SNPs based on their number of minor allele copies present in either population. The following data frame shows the number of SNPs for each bin class:

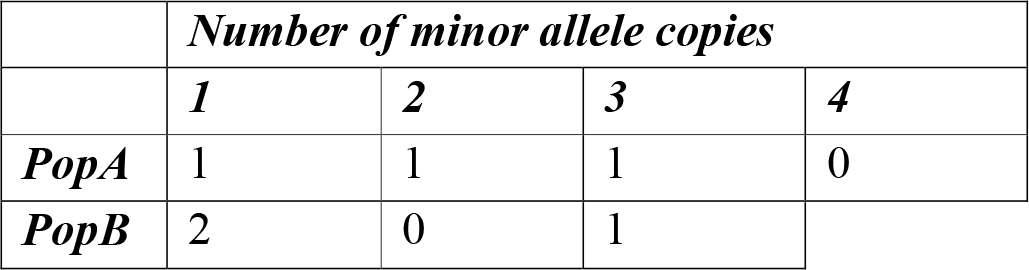

Hence the site frequency spectra are:

PopA: 1,1,1,0
PopB: 2,0,1

Note that the row sums are equal to the number of segregating sites within either population. The length of the vectors is equal to the number of individuals within either population.

## Notes

https://github.com/mennodejong1986/SambaR

### Competing Interest Statement

The authors have declared no competing interest.

## References

Abel, G. J. (2019). migest: Methods for the Indirect Estimation of Bilateral Migration. https://CRAN.R-project.org/package=migest

Adler, D., & Kelly, S. T. (2019). vioplot: Violin plot. https://github.com/TomKellyGenetics/vioplot

Alexander, D. H., Novembre, J., & Lange, K. (2009). Fast model-based estimation of ancestry in unrelated individuals. Genome Research, 19(9), 1655–1664. https://doi.org/10.1101/gr.094052.109

Auguie, B. (2017). gridExtra: Miscellaneous Functions for “Grid” Graphics. https://CRAN.R-project.org/package=gridExtra

Baudouin & Cornuet, 2004, Analytical Bayesian Approach for Assigning Individuals to Populations. Journal of Heredity, 95(3), 217–224.

Biscarini, F., Cozzi, P., Gaspa, G. and Marras, G. (2018). detectRUNS: Detect runs of homozygosity and runs of heterozygosity in diploid genomes. CRAN (The Comprehensive R Archive Network)

Bosse, M., Spurgin, L. G., Laine, V. N., Cole, E. F., Firth, J. A., Gienapp, P., Gosler, A. G., McMahon, K., Poissant, J., Verhagen, I., Groenen, M. A. M., Van Oers, K., Sheldon, B. C., Visser, M. E., Slate, J. (2017). Recent natural selection causes adaptive evolution of an avian polygenic trait. Science, 358(6361), 365–368

Caye, K., Jay, F., Michel, O., & François, O. (2018). Fast inference of individual admixture coefficients using geographic data. The Annals of Applied Statistics, 12(1), 586–608. https://doi.org/10.1214/17-AOAS1106

Chang, C. C., Chow, C. C., Tellier, L. C., Vattikuti, S., Purcell, S. M., & Lee, J. J. (2015). Second-generation PLINK: Rising to the challenge of larger and richer datasets. GigaScience, 4, 7. https://doi.org/10.1186/s13742-015-0047-8

Chen, H. (2018). VennDiagram: Generate High-Resolution Venn and Euler Plots. https://CRAN.R-project.org/package=VennDiagram

Cockerham and Weir, 1987, Correlations, descent measures: Drift with migration and mutation

De Jong, M., Li, Z., Qin, Y., Quemere, E., Baker, K., Wang, W., Hoelzel, A.R. (2020). Demography and adaptation promoting evolutionary transitions in a mammalian genus diversifying during the Pleistocene. Molecular Ecology. https://doi.org/10.1111/mec.15450

Durand, E. Y., Patterson, N., Reich, D., Slatkin, M. (2011). Testing for ancient admixture between closely related populations. Molecular Biology and Evolution, 28(8), 2239–2252.

Flanagan, S. P., & Jones, A. G. (2018). fsthet: Fst-Heterozygosity Smoothed Quantiles. https://CRAN.R-project.org/package=fsthet

Foll, M., & Gaggiotti, O. (2008). A Genome-Scan Method to Identify Selected Loci Appropriate for Both Dominant and Codominant Markers: A Bayesian Perspective. Genetics, 180(2), 977–993. https://doi.org/10.1534/genetics.108.092221

Frichot, E., & François, O. (2015). LEA: An R package for landscape and ecological association studies. Methods in Ecology and Evolution, 6(8), 925–929. https://doi.org/10.1111/2041-210X.12382

Fung, T. & Keenan, K. (2014). Confidence intervals for population allele frequencies: the general case of sampling from a finite diploid population of any size. PloS One, 9(1). https://doi.org/10.1371/journal.pone.0085925

Funk, W. C., Lovich, R. E., Hohenlohe, P. A., Hofman, C. A., Morrison, S. A., Scott Sillett, T., Ghalambor, C. K., Maldonado, J. E., Rick, T. C., Day, M. D., Polato, N. R., Fitzpatrick, S. W., Coonan, T. J., Crooks, K. R., Dillon, A., Garcelon, D. K., King, J. L., Boser, C. L., Gould, N., Andelt, W. F. (2016). Adaptive divergence despite strong genetic drift: genomic analysis of the evolutionary mechanisms causing genetic differentiation in the island fox (Urocyon littoralis). Molecular Ecology. 25(10)

Gaunt, T. R., Rodríguez, S., & Day, I. N. (2007). Cubic exact solutions for the estimation of pairwise haplotype frequencies: Implications for linkage disequilibrium analyses and a web tool “CubeX.” BMC Bioinformatics, 8, 428. https://doi.org/10.1186/1471-2105-8-428

Gel, B., Serra, E. (2017). karyoploteR: an R / Bioconductor package to plot customizable genomes displaying arbitrary data. Bioinformatics, 33(9), 3088–3090

Gerritsen, H. (2018). mapplots: Data Visualisation on Maps. https://CRAN.R-project.org/package=mapplots

Gu, Z., Gu, L., Eils, R., Schlesner, M., & Brors, B. (2014). Circlize implements and enhances circular visualization in R. Bioinformatics, 30(19), 2811–2812.

Heppenheimer, E., Brzeski, K. E., Hinton, J. W., Patterson, B. R., Rutledge, L. Y., DeCandia, A. L., Wheeldon, T., Fain, S. R., Hohenlohe, P. A., Kays, R., White, B. N., Chamberlain, M. J., vonHoldt, B. M. (2018). High genomic diversity and candidate genes associated with a range expansion in eastern coyote (*Canis latrans*) populations. Ecology and Evolution. https://doi.org/10.1002/ece3.4688

Hijmans, R. J. (2019). raster: Geographic Data Analysis and Modeling. https://CRAN.R-project.org/package=raster

Hudson, R.R., Slatkin, M., Maddison, W.P. (1992), Estimation of levels of gene flow from DNA sequence data. Genetics, 132(2), 583–589

Jombart, T. (2008). adegenet: An R package for the multivariate analysis of genetic markers. Bioinformatics, 24, 1403–1405. https://doi.org/10.1093/bioinformatics/btn129

Jombart, T., Devillard, S., & Balloux, F. (2010). Discriminat analysis of principal components: a new method for the analysis of genetically structured populations. BMC Genetics 11(94)

Jombart, T., & Ahmed, I. (2011). adegenet 1.3-1: New tools for the analysis of genome-wide SNP data. Bioinformatics. https://doi.org/10.1093/bioinformatics/btr521

Kamvar, Z. N., Tabima, J. F., & Grunwald, N. J. (2014). *Poppr*: An R package for genetic analysis of populations with clonal, partially clonal, and/or sexual reproduction. PeerJ, 2, e281. https://doi.org/10.7717/peerj.281

Kardos, M., Luikart, G., Allendorf, F.W. (2015). Measuring individual inbreeding in the age of genomics: marker-based measures are better than pedigrees. Heredity, 115, 63–72.

Kassambara, A., & Mundt, F. (2019). factoextra: Extract and Visualize the Results of Multivariate Data Analyses. https://CRAN.R-project.org/package=factoextra

Lê, S., Josse, J., & Husson, F. (2008). FactoMineR: An R Package for Multivariate Analysis. Journal of Statistical Software, 25(1), 1–18. https://doi.org/10.18637/jss.v025.i01

Ligges, U., & Mächler, M. (2003). Scatterplot3d—An R Package for Visualizing Multivariate Data. Journal of Statistical Software, 8(11), 1–20.

Liu, S., Lorenzon, E. D., Fumagalli, M., Li, B., Harris, K., Xiong, Z., Zhou, L., Korneliussen, T. S., Somel, M., Babbitt, C., Wray, G., Li, J., He, W., Wang, Z., Fu, W., Xiang, X., Morgan, C. C., Doherty, A., O’Connell, M. J., McInerney, J. O., Born, E. W., Dalén, L., Dietz, R., Orlando, L., Sonne, C., Zhang, G., Nielsen, R., Willerslev, E., Wang, J. (2014). Population Genomics Reveal Recent Speciation and Rapid Evolutionary Adaptation in Polar Bear. Cell, 157(4), 785–794

Liu, X., & Fu, Y.-X. (2015). Exploring Population Size Changes Using SNP Frequency Spectra. Nature Genetics, 47(5), 555–559. https://doi.org/10.1038/ng.3254

Luu, K., Bazin, E., & Blum, M. G. B. (2017). pcadapt: An R package to perform genome scans for selection based on principal component analysis. Molecular Ecology Resources, 17(1), 67–77. https://doi.org/10.1111/1755-0998.12592

Luu, K., Blum, M., & Prive, F. (2019). pcadapt: Fast Principal Component Analysis for Outlier Detection. https://CRAN.R-project.org/package=pcadapt

Murrell, P. (2005). R Graphics. Chapman & Hall/CRC Press.

Murrell, Paul, & Wen, Z. (2019). gridGraphics: Redraw Base Graphics Using “grid” Graphics. https://CRAN.R-project.org/package=gridGraphics

Mussmann, S. M., Douglas, M. R., Chafin, T. K., & Douglas, M. E. (2019). BA3-SNPs: Contemporary migration reconfigured in BayesAss for next-generation sequence data. Methods in Ecology and Evolution, 10(10), 1808–1813. https://doi.org/10.1111/2041-210X.13252

Nei, Chakravarti, Tateno, 1977, Mean and variance of Fst in a finite number of incompletely isolated populations

Neuwirth, E. (2014). RColorBrewer: ColorBrewer Palettes. https://CRAN.R-project.org/package=RColorBrewer

Paradis, E., & Schliep, K. (2018). ape 5.0: An environment for modern phylogenetics and evolutionary analyses in R. Bioinformatics, 35, 526–528.

Peatkau, D., Calvert, W., Stirling, I., Strobeck, C. (1995). Microsatellite analysis of population structure in Canadian polar bears. Molecular Ecology. https://doi.org/10.1111/1j.1365-294X.1995.tb00227.x

Peuchmaille, 2016, The programs structure does not reliably recover the correct population structure when sampling is uneven. Mol Ecol Resour, 16: 608–627

Pembleton, L. W., Cogan, N. O. I., & Forster, J. W. (2013). StAMPP: an R package for calculation of genetic differentiation and structure of mixed-ploidy level populations. Molecular Ecology Resources, 13, 946–952. https://doi.org/10.1111/1755-0998.12129

Purcell, S., Neale, B., Todd-Brown, K., Thomas, L., Ferreira, M. A. R., Bender, D., Maller, J., Sklar, P., de Bakker, P. I. W., Daly, M. J., & Sham, P. C. (2007). PLINK: A Tool Set for Whole-Genome Association and Population-Based Linkage Analyses. The American Journal of Human Genetics, 81(3), 559–575. https://doi.org/10.1086/519795

R Core Team. (2019). R: A Language and Environment for Statistical Computing. R Foundation for Statistical Computing. https://www.R-project.org/

Soetaert, K. (2017). Plot3D: Plotting Multi-Dimensional Data. https://CRAN.R-project.org/package=plot3D

South, A. (2011). rworldmap: A New R package for Mapping Global Data. The R Journal, 3, 35–43. https://doi.org/10.32614/RJ-2011-006

Storey, J. D., Bass, A. J., Dabney, A., & Robinson, D. (2019). qvalue: Q-value estimation for false discovery rate control. http://github.com/jdstorey/qvalue

Viengkone, M., Derocher, A. E., Richardon, E. S., Malenfant, R. M., Miller, J. M. Obbard, M. E., Dyck, M. G., Lunn, N. J., Sahanatien, V., Davis, C. S. (2017). Assessing polar bear (*Ursus maritimus*) population structure in the Hudson Bay region using SNPs. Ecology and Evolution, 6(23), 8474–8484.

Von Thaden, A., Nowak, C., Tiesmeyer, A., Reiners, T. E., Alves, P. C., Lyons, L. A., Mattucci, F., Randi, E., Cragnoloni, M., Galian, J., Hegyeli, Z., Kitchener, A. C., Lambinet, C., Lucas, J. M., Mölich, T., Ramos, L., Schockert, V., Cocchiararo, B. 2020, Applying genomic data in wildlife monitoring: Development guidelines for genotyping degraded samples with reduced single nucleotide polymorphism panels. Molecular Ecology Resources. https://doi.org/10.1111/1755-0998.13136

Waples, R. K., Albrechtsen, A., Moltke, I. (2018). Allele frequency-free inference of close familial relationships from genotypes or low depth sequencing data. Molecular Ecology, 28

Warnes, G. R., Bolker, B., Bonebakker, L., Gentleman, R., Liaw, W. H. A., Lumley, T., Maechler, M., Magnusson, A., Moeller, S., Schwartz, M., & Venables, B. (2019). gplots: Various R Programming Tools for Plotting Data. https://CRAN.R-project.org/package=gplots

Ward, B. J & Van Oosterhout, C. (2016). HYBRIDCHECK: software for the rapid detection, visualization and dating of recombination regions in genome sequence data. Molecular Ecology Resources, 16(2): 534–539

Weir and Cockerham, 1984, Estimating F-statistics for the analysis of population structure. Evolution, 38(6), 1558–1370.

Weldenegodguad, M., Pokharel, K., Ming, Y., Honkatukia, M., Peippo, J., Reilas, T., Roed, K. H., Kantanen, J. (2019). Genome sequence and comparative analysis of reindeer (*Rangifer tarandus*) in northern Eurasia. Scientific Reports, 10, https://doi.org//10.1038/s41598-020-65487-y

Whitlock, M. C., & Lotterhos, K. (2014). OutFLANK: Fst outliers with trimming.

Whitlock, M. C., & Lotterhos, K. E. (2015). Reliable Detection of Loci Responsible for Local Adaptation: Inference of a Null Model through Trimming the Distribution of F(ST). The American Naturalist, 186 Suppl 1, S24–36. https://doi.org/10.1086/682949

Wickham, H. (2011). The Split-Apply-Combine Strategy for Data Analysis. Journal of Statistical Software, 40(1), 1–29.

Wickham, H., & Seidel, D. (2019). scales: Scale Functions for Visualization. https://CRAN.R-project.org/package=scales

Wright, S. (1943), Isolation by distance. Genetics, 28 (2), 114–138.

Yang, J., Hong Lee, S., Goddard, M. E., Visscher, P. M. (2011). GCTA: A Tool for Genome-wide Complex Trait Analysis. Am J Hum Genet. 2011, 88(1), 76–82

Zeileis, A., Fisher, J. C., Hornik, K., Ihaka, R., McWhite, C. D., Murrell, P., Stauffer, R., & Wilke, C. O. (2019). colorspace: A Toolbox for Manipulating and Assessing Colors and Palettes (ArXiv 1903.06490). arXiv.org E-Print Archive. http://arxiv.org/abs/1903.06490

Zeileis, A., & Grothendieck, G. (2005). zoo: S3 Infrastructure for Regular and Irregular Time Series. Journal of Statistical Software, 14(6), 1–27. https://doi.org/10.18637/jss.v014.i06

